# Arabidopsis Transcription Factor WRKY45 Confers Cadmium Tolerance via Activating *PCS1* and *PCS2* Expression

**DOI:** 10.1101/2023.07.17.549240

**Authors:** Fangjian Li, Yaru Deng, Yan Liu, Cuishan Mai, Yun Xu, Jiarui Wu, Xinni Zheng, Cuiyue Liang, Jinxiang Wang

**Affiliations:** Root Biology Center, South China Agricultural University, Guangzhou 510642, P.R. China; College of Natural Resources and Environment, South China Agricultural University, Guangzhou 510642, P. R. China; Key Laboratory of Agricultural and Rural pollution Control and Environmental Safety in Guangdong Province, Guangzhou 510642, P. R. China; State Key Laboratory for Conservation and Utilization of Subtropical Agro-bioresources, South China Agricultural University, Guangzhou 510642, P. R. China

**Keywords:** Cd stress, WRKY45, PCS1, PCS2, Arabidopsis, PC

## Abstract

Cadmium (Cd) has long been recognized as toxic pollutant to crops worldwide. The biosynthesis of glutathione-dependent phytochelatin plays crucial roles in the detoxification of Cd in plants. However, its regulatory mechanism remains elusive. Here, we revealed that Arabidopsis transcription factor WRKY45 confers Cd tolerance via promoting the expression of PC synthesis-related genes PCS1 and PCS2, respectively. Firstly, we found that Cd stress induces the transcript levels of WRKY45 and its protein abundance. Accordingly, in contrast to wild type Col-0, the increased sensitivity to Cd is observed in *wrky45* mutant, while overexpressing WRKY45 plants are more tolerant to Cd. Secondly, quantitative real-time PCR revealed that the expression of AtPCS1 and AtPCS2 is stimulated in overexpressing WRKY45 plants, but decreased in wrky45 mutant. Thirdly, WRKY45 promotes the expression of PCS1 and PCS2, electrophoresis mobility shift assay analysis uncovered that WRKY45 directly bind to the W-box cis-element of PCS2 promoter. Lastly, the overexpression of WRKY45 in Col-0 leads to more accumulation of PCs in Arabidopsis, and the overexpression of PCS1 or PCS2 in *wrky45* mutant plants rescues the phenotypes induced by Cd stress. In conclusion, our results show that AtWRKY45 positively regulate Cd tolerance in Arabidopsis via activating PCS1 and PCS2 expression.

**Environmental implication:** Accumulation of cadmium (Cd) in soils poses a threat to crop productivity and food safety. It has been revealed that phytochelatin (PC) plays an essential role in plants to alleviate Cd toxicity, yet the regulatory mechanisms governing its expression remain unclear. We have demonstrated that the Arabidopsis transcription factor *WRKY45* directly activates the expression of *PCS1* and *PCS2*, which encode PC synthase, thereby increasing the content of PC and enhancing Arabidopsis tolerance to Cd stress. These findings offer insights into precise regulation strategies for crop Cd tolerance via modulation of *WRKY45* homologue in crops.

## 1. Introduction

Heavy metal cadmium (Cd) is highly toxic to both animals and plants (Huang et al., 2017; Reeves and Chaney, 2008). Due to the rapid increase of soil Cd content caused by human activities, such as mining, smelting, sewage irrigation and fertilization, Cd has become one of the most dangerous environmental pollutants especially for the important crop rice in Asia (Fu et al., 2021; Huang et al., 2023; Tong et al., 2023; Zhao et al., 2023). As revealed, more than 7.0% of the soils in China has excessive Cd content, so it is urgent to control the soil contaminated by Cd (Meng et al., 2022). Using genetic technology to select Cd-tolerant crop varieties for bioremediation is a promising technology for the treatment of Cd-contaminated soils because of its environmental friendliness and low cost (Clemens, 2019; DalCorso et al., 2008; Grant et al., 2008; Wang et al., 2019; Wu et al., 2023). Therefore, it is of great significance to understand the tolerance mechanism of plants to Cd toxicity.

Plants have evolved elegant mechanisms to detoxify and tolerate Cd stress (Tang et al., 2023; Zhao et al., 2022). These mechanisms include the synthesis of small molecular substances that chelate Cd, isolating the complex of Cd with glutathione (GSH) or Phytochelatin (PC) in vacuoles, reducing the absorption of Cd, excreting intracellular ions, arresting Cd on cell wall, and the interaction between plants and root microorganisms to reduce Cd stress (Clemens, 2001; Dang et al., 2022; Kim et al., 2007; Peng et al., 2017; Zhu et al., 2020).

Being a kind of small molecular substance, PC can chelate Cd efficiently by using GSH as precursor and catalyzed by PC synthetase (Tang et al., 2023). The intracellular PC-Cd complexes is isolated after entering the vacuole through the tonoplast carrier ABCC1/2 (Zhao et al., 2022). Some studies have shown that PCs also detoxify other toxic metals such as Hg and Pb (Cohen et al., 1998; Fischer et al., 2014). In *Arabidopsis thaliana*, transcription factors *ZAT6* and *MYB4* directly or indirectly positively regulate the transcription of several genes in the GSH and PC biosynthesis pathway, thus affecting the tolerance to Cd (Agarwal et al., 2020; Chen et al., 2016). In Arabidopsis, *GSH1* and *GSH2* serve as pivotal enzymes for the production of GSH, while PCS1 and PCS2 facilitate the biosynthesis of PC (Cohen et al., 1998; Fischer et al., 2014). Accordingly, the mutation of *GSH1*, *GSH2*, *PCS1* or *PCS2* results in more sensitive to Cd stress in Arabidopsis. As reported, under Cd stress, *WRKY12* represses the expression of *GSH1* (Han et al., 2019). However, the regulation of *PCS1* and *PCS2* under Cd stress is unclear.

WRKY transcription factor is one of the plant-specific gene families, which regulate plant growth and development, and adaptations to biotic and abiotic stresses (Dong et al., 2003; Eulgem et al., 2000). To date, a total of 74 *WRKY* members have been identified in the reference plant *Arabidopsis thaliana* genome (Ulker and Somssich, 2004). Many studies have demonstrated the crucial roles of WRKY family members in plant responses to diverse environmental stresses, including nutrient deficiencies, heavy metal toxicity, high salinity stress, hypoxia response, as well as biotic challenges from diseases and pests (Chen et al., 2017; Devaiah et al., 2007; Han et al., 2019; Hu et al., 2012; Li et al., 2011; Li et al., 2013; Scarpeci et al., 2013; Sheng et al., 2019; Su et al., 2015; Sun et al., 2020; Wang et al., 2018; Zheng et al., 2006). *WRKY45* has been shown to be involved in the regulation of dark-induced leaf senescence in *Arabidopsis thaliana* (Barros et al., 2022; Chen et al., 2017), phosphorus uptake (Wang et al., 2014), and immune resistance in rice via regulating the expression of downstream genes (Goto et al., 2015; Shimono et al., 2012). *WRKY45* directly regulate the expression of phosphate transporter *Pht1;1* in Arabidopsis via binding the W-box *cis*-element (TTGACC/T) (Wang et al., 2014). However, it is not clear whether *WRKY45* plays a role in heavy metal detoxification in *Arabidopsis thaliana*.

In this study, we found that *WRKY45* is involved in regulating the important role of plants in Cd stress. The expression of *WRKY45* gene and protein level are enhanced under Cd stress. Consistent with this, the promoter activities of *WRKY45* are also increased by Cd stress both in roots and leaves, respectively. The overexpression of *WRKY45* leads to the increase of PC content and the enhancement of tolerance to Cd, while the loss of *WRKY45* function leads to the decrease of PC content, thus resulting in the enhanced sensitivity to Cd. Moreover, Dual-LUC analysis confirmed that *WRKY45* directly activates the expression of *PCS1* and *PCS2*, which encode PC synthase. Additionally, an EMSA experiment revealed that WRKY45 binds to the W-box cis-element in the promoter region of *PCS2*. Of note, the overexpression of *PCS1* or *PCS2* in *wrky45* mutant can restore the tolerance of *wrky45* to Cd stress, indicating that *PCS1* and *PCS2* function downstream of *WRKY45*.

## 2. Materials and methods

### 2.1 Growth conditions

The *Arabidopsis thaliana* (L.) Columbia (Col-0) was used in this study. The *wrky45* mutant (GK-684G12), WRKY45-OX-15, HA-WRKY45 and WRKY45-GUS line (Chen et al., 2017) seeds were kindly provided by Dr. Diqiu Yu (Key Laboratory of Tropical Plant Resources and Sustainable Use, Xishuangbanna Tropical Botanical Garden, Chinese Academy of Sciences, Kunming, Yunnan 650223, China). The seeds of PC-biosynthesis related mutants *cad1-3*(CS68125) and *pcs2* (SAIL_1254_H08) were purchased from Arabidopsis Biological Resource Center (www.arabidopsis.org).

The seeds were surface sterilized with 75% alcohol for 3 minutes, then soaked in 95% alcohol for 30 seconds and dried. After sterilization, the seeds were sowed on half strength Murashige and Skoog (1/2 MS) media with 0.8% sucrose and 1% Phytagel (Sigma-Aldrich) and kept the plates in the dark at 4 °C for 2 days for stratification treatment. The seeds are then placed in a growth chamber with a thermocycle of 22 ℃/20 ℃ and a photoperiod of 16 h light/8 h dark, with a light intensity of 100 μmol^-2^s^-1^ as described (Ou et al., 2022). For Cd treatment, CdCl_2_ was directly added to the growth media as indicated.

### 2.2 Plasmid construction and plant transformation

The plant overexpression binary vector pCAMBIA1300 containing constitutive 35S promoter was used to construct the overexpression vector. The sequence information of all genes comes from TAIR (Arabidopsis thaliana information resource www.arabidopsis.org). The full-length cDNA of each gene was amplified from the Arabidopsis cDNA. The amplification products of *PCS1* and *PCS2* were cloned into pCAMBIA1300 digested with SalI and PstI, respectively.

PCS1-OX and PCS2-OX were introduced into *Agrobacterium tumefaciens* strain GV3101 by freeze-thaw method, and *wrky45* mutants were transformed by flower dip method (Clough and Bent, 1998). The transgenic lines used in this study are all T_3_ homozygous plants with single copy insertion via Chi-test. In addition, the *wrky45* mutant (GK-684G12) plants were transformed with plasmids that contains *PCS1* or *PCS2* open reading frame driven by 35S promoter, respectively to create *wrky45* PCS1-OX or *wrky45* PCS2-OX lines. The sequences of primer pairs used for vector construction are shown in Supplementary Table S1.

### 2.2 GUS activity detection

The transgenic Arabidopsis seeds germinated on half strength MS agar medium and transferred to half-strength MS agar media with or without 75 μM CdCl_2_ 3 days after germination. Then GUS staining was performed at 6^th^ hour Cd treatment after seedling transfer as described (Tang et al., 2022).

### 2.3 Measurement of plant primary root length and fresh weight

After capturing images of the sample with digital camera Nikon D300s, the primary root length was quantified through ImageJ (Ou et al., 2022). Statistical analysis was performed based on three biological experiments.

### 2.4 RNA extraction and quantitative RT -PCR analysis

The total RNA from *Arabidopsis* seedlings was extracted and purified according to the previous methods (Zhu et al., 2020), and the purity was evaluated by A260/A280 ratio using NanoDrop spectrophotometer (Thermo, USA). The total RNA was transcribed back into cDNA (ABclonal, China, RK20429 https://abclonal.com.cn/) using a reverse transcription kit. The cDNA was used for quantitative RT-PCR analysis of SYBR Green monitoring on the 7500 real-time PCR system (Thermo, USA). The primer pairs used for quantitative real time PCR (qRT-PCR) analysis of *WRKY45*, *GSH1*, *GSH2*, *PCS1*, *PCS2* and housekeeping gene *ACTIN2* (*AT3G18780*) were list in Supplementary Table S1.

### 2.5 Analysis of cadmium and iron contents

Seedlings were grown on full hydroponic nutrient solutions for 2 weeks, and then treated with 20 µM CdCl_2_ for 24 hours (h). Thus, the plants were sampled and the contents of Cd were analyzed according to the described method (Lee et al., 2007). At the same time, the content of iron was determined the digested samples were analyzed by atomic absorption spectrometer (Solaar M6, ThermoFisher, Waltham, USA). The Cd accumulation and translocation factor are calculated according to the method described (Bose and Bhattacharyya, 2008; Meng et al., 2022).

### 2.6 Dual-Luciferase Assay

Dual-Luciferase experiments were carried out as described (Liu et al., 2008), and the procedure was modified. *PCS1* and *PCS2* promoters were cloned into pGreen-0800LUC vector as reporter vectors (PCS1Pro-LUC and PCS2Pro-LUC). The effector 35S:WRKY45 was constructed by using pGreen62-SK vector. As mentioned above, both the reporter plasmid and the effector plasmid were simultaneously transferred into *Nicotiana benthamiana* leaves for LUC detection. Double LUC was detected by double luciferase reporter gene detection kit (YeSen, China https://www.yeasen.com/). LUC and REN signals are detected using a modular microboard multimode reader (Turner Biossystem). Three biological repeat sequences were measured in each sample and similar results were obtained.

### 2.7 Protein extraction and immunoblot analysis

Total protein was extracted as described before (Tang et al., 2022). *Arabidopsis* plants were ground in liquid N_2_ and extracted with extraction buffer (50 mM Tris, pH 7.5; 150 mM NaCl; 5 mM EDTA; 2 mM DTT; 10% glycerol; 1 mM PMSF). After centrifugation at 12 000 rpm for 10 min at 4°C, supernatants were loaded on 10% SDS-PAGE for Western blotting using anti-HA (M20003) or anti-ACTIN antibody (M20009) (Abmart, http://www.ab-mart.com.cn/, China).

### 2.8 Electrophoresis mobility shift assay (EMSA)

EMSA was carried out according to the manufacturer’s protocol using the EMSA/Gel-Shift kit (GS009, Beyotime, https://www.beyotime.com/index.htm, China). The whole coding sequence of *WRKY45* was amplified by primer GST_WRKY45_F and GST_WRKY45_R, respectively. The amplification product of *WRKY45* was cloned into pGEX-6P-3 vector (GE Healthcare, USA) that was cleaved with BamHI. The expression vector was introduced into *E. coli* BL21 strain. The recombinant protein was extracted with BeverBeadsTM GSH magnetic beads (70601-K10, Beaver Nano, https://www.beaverbio.com/, China). Oligonucleotide probes (PCS1.EMSA-biotinR and PCS2.EMSA-biotinR) were synthesized and labeled with biotin at the 5’end of the sense chain. All the primer sequences mentioned above are shown in Supplementary Table S1.

### 2.9 Analysis of the contents of PC, NPT, GSH, and Chlorophyll

The two-week-old seedlings hydroponic cultured were treated with 20 µM CdCl_2_ for 24 hours, then the contents of PC, NPT, GSH were analyzed and determined as described (Chen et al., 2015).

The seedlings were cultured on half-strength MS media with or without 75 μ M CdCl_2_ for 10 days, and then the chlorophyll content was analyzed based on published method (Nam et al., 2021).

## 3. Results

### 3.1 The transcripts of *AtWRKY45* and protein level of WRKY45 are enhanced under Cd stress

As documented, the members of the *WRKY* family of *Arabidopsis thaliana* such as *WRKY12*, *WRKY13* and *WRKY33* are involved in the regulation of Cd stress (Han et al., 2019; Sheng et al., 2019; Zhang et al., 2023). We speculate that other *WRKY* genes may also respond to Cd stress. To test this hypothesis, we screened some up-regulated *WRKY* genes in *Arabidopsis* under Cd stress in the transcriptome database (Kong et al., 2018) and found that *AtWRKY45* was reported to be upregulated by Cd. Thus, we used quantitative real time PCR (qRT-PCR) to explore the transcripts of *AtWRKY45* to Cd. As shown, the transcript level of *WRKY45* in *Arabidopsis thaliana* roots or leaves is promoted by Cd stress, respectively. In contrast to the control (without Cd application), the expression of *WRKY45* peaks at the 6th hour after Cd stress and then decreases over time in both roots (Fig.1A) and leaves (Fig. 1B). When treated with 6-h Cd, the abundance of *WRKY45* is upregulated by 3.5 folds in leaves (Fig.1A), and 3.4 folds in roots (Fig.1B), respectively. Our results verified that *AtWRKY45* is induced by Cd stress.

**Fig. 1.**
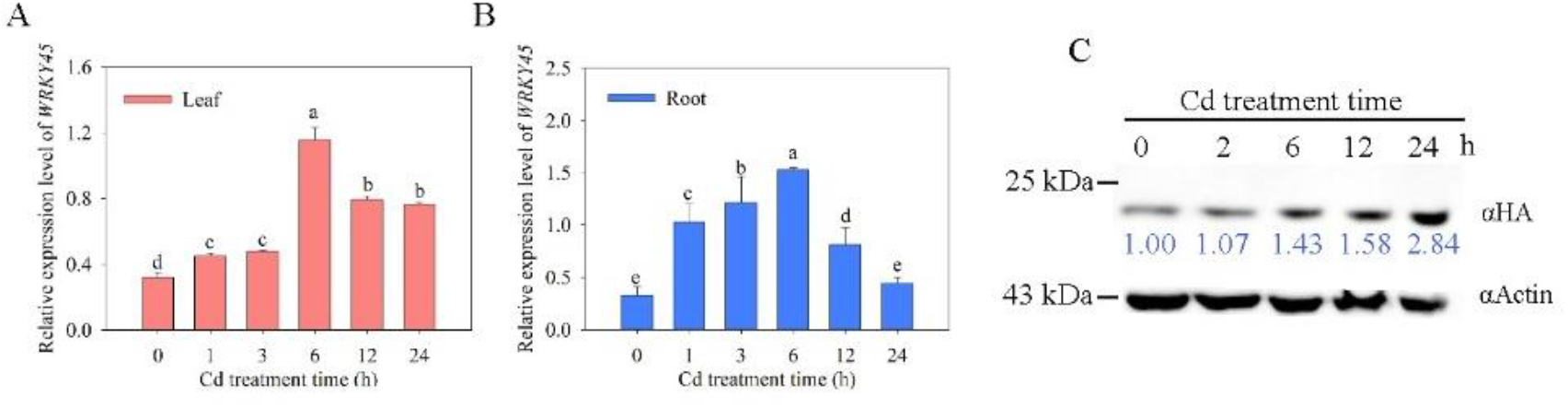
The relative expression level of *WRKY45* in Arabidopsis roots (A) and leaves. (B) qRT-PCR analysis of transcript levels of *WRKY45* Arabidopsis leaves and roots, respectively, at different time point after Cd stress. The relative expression level of *WRKY45* was normalized against *ACTIN2* (*AT3G18780*). Data represent means ± SD (n = 4) from four independent experiments. Different letters indicate significant difference at P<0.05 (one-way ANOVA with Turkey’s test). (C) Immunoblot analysis of HA-WRKY45 protein in the plant of HA-WRKY45 line in response to Cd stress. Plants were grown on nutrient solution for 2 weeks, and after treated with 20 µM CdCl_2_ for 0, 2, 6,12, 24 h, respectively, then total protein was extracted and analyzed using anti-HA or anti-ACTIN antibody. Relative amounts of proteins were determined by densitometry using ImageJ software, ACTIN was used as the loading control.

Then we employed the proWRKY45: GUS line plants as reported and assessed the activities of the *WRKY45* promoter under Cd stress. Under control conditions (no Cd application), we observed the activities of the *WRKY45* promoter in roots, stems, and leaves. Notably, under Cd stress conditions, the activities of the *WRKY45* promoter in both roots and leaves are significantly increased (Fig. 2). This results further support that *WRKY45* positively respond to Cd.

**Fig. 2.**
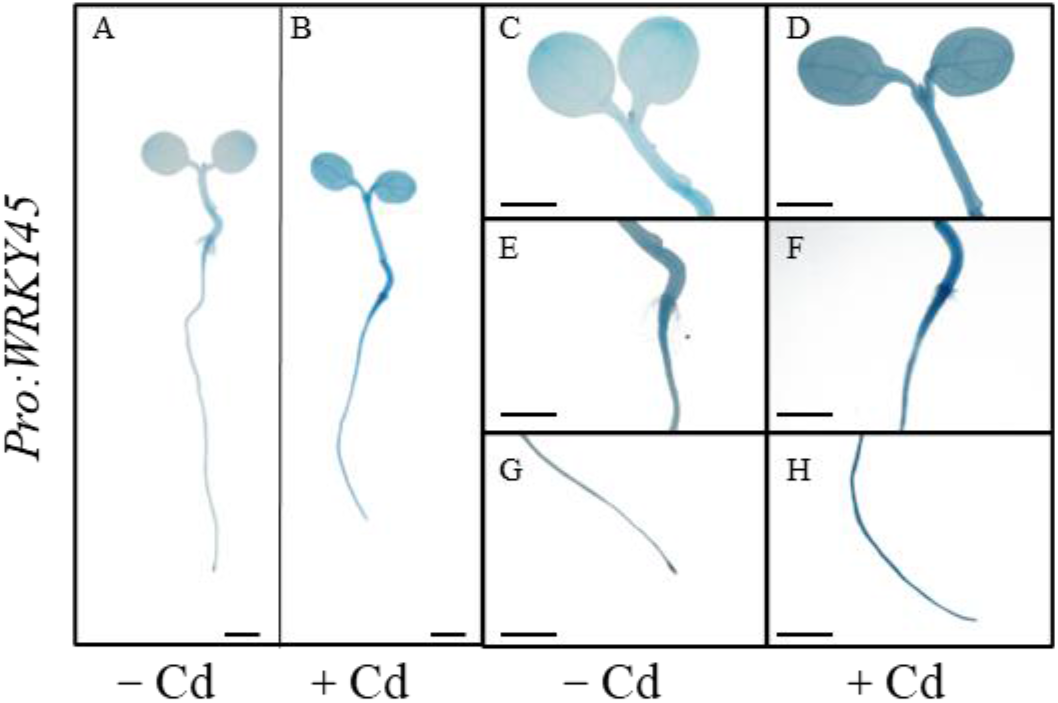
The activities of *WRKY45* promoter are stimulated by Cd stress. Histochemical GUS staining was performed in the proWRKY45: GUS seedlings. (A, B) Arabidopsis seeds were germinated on basal agar medium, transferred to solid agar media with or without 75 μM CdCl_2_ at 3 d after germination, and then analyzed at 6 h after seedling transfer, scale bar, 1 mm; (C, D); cotyledon, scale bar, 2 mm; (E, F) root connection part, scale bar, 2 mm; (G, H) primary root, scale bar, 2 mm.

Next, we utilized the WRKY45: HA transgenic line (Chen et al., 2017) to investigate the abundance of WRKY45 via Western Blot. As demonstrated in Fig. 1C, the level of AtWRKY45 increases with prolonged exposure to Cd (Fig. 1C). The levels of WRKY45 gradually increases within a 24-hour period of Cd stress and are found to be 1.43, 1.58, and 2.8 times higher at Cd-stressed 6 h, 12 h, 24 h than the control. This verified that Cd stress increases the abundance of WRKY45.

### 3.2 WRKY45 positively modulates cadmium tolerance

To decipher the physiological functions of *WRKY45* in Cd stress, we thus used the previously reported *wrky45* mutant and the WRKY45-OX-15 overexpressing line (Chen et al., 2017). We verified that the *wrky45* mutant results from the inserted T-DNA into the 5 ’untranslated region of the *WRKY45* gene (Fig. S1A), and genomic PCR analysis confirmed the T-DNA insertion (Fig. S1B). Of note, the expression level of *WRKY45* in *wrky45* seedlings is approximate 20% of that in wild type Col-0, regardless of whether Cd was applied or not. Conversely, the transcripts of *WRKY45* in WRKY45-OX-15 seedlings is ten-fold higher than that in Col-0. (Fig. S1C). These results confirm that the T-DNA insertion line functions as a knock-down line of *WRKY45*, while *WRKY45-OX-15* serves as the overexpressing line. This is consistent with previous study (Chen et al., 2017). We therefore employed the two lines in subsequent experiments.

Compared to Col-0, no significant growth and developmental defects were observed in both *wrky45* and WRKY45-OX-15 lines during the seedling stage under half-strength media. However, when exposed to Cd, the overexpressing WRKY45 line plants exhibit more Cd tolerance than Col-0, while the *wrky45* mutant plants are more sensitive to Cd than Col-0 (Fig. 3A). We thus quantified primary root length (Fig. 3B), fresh weight (Fig. 3C), and chlorophyll content (Fig. 3D). Under 0 μM CdCl_2_ conditions, no difference can be found among Col-0, *wrky45* and WRKY45-OX-15 plants. But stressed with 50 μM CdCl_2_ and 75 μM CdCl_2_, the *wrky45* mutant plants exhibited a reduction in primary root length by 18.4% and 27.0%, respectively, while the WRKY45-OX-15 plants showed an increase of 7.9% and 18.2%, respectively (P <0.05, Fig.3B). The fresh weight of the *wrky45* plants decreased by 27.3% and 30.3%, respectively, under these conditions, whereas that of the overexpressing WRKY45-OX-15 plants increased by 7.9% and 18.2%, respectively (P <0 .05, Fig.3C).

**Fig. 3.**
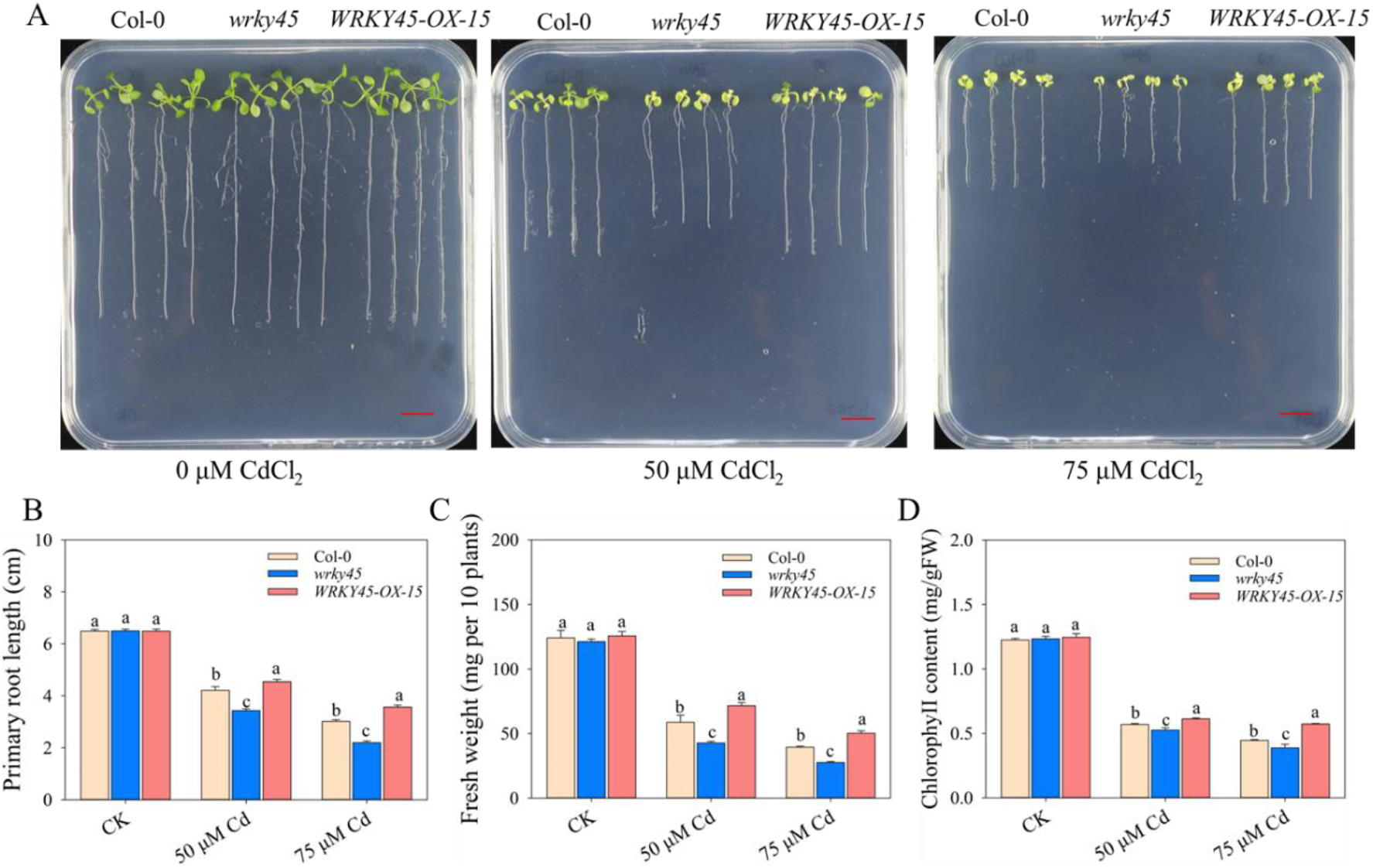
The *wrky45* mutants are sensitive to Cd stress; and the WRKY45-OX genotypes are tolerant to Cd stress. (A) The loss of *WRKY45* function results in more sensitive to Cd stress. Seedlings were grown on half-strength MS media for 2 days then treated with 0, 50, or 75 µM CdCl_2_ for 10 days. Scale bar, 1 cm. (B-D) Primary root length (B), fresh weight (C) and chlorophyll content (D) of wild-type (Col-0), *wrky45* and WRKY45-OX plants described in A. Data present the means ± SE (n = 4). Different letters indicate significant difference at P < 0.05 level (one-way ANOVA with Turkey’s test).

As reported, application of Cd inhibits iron uptake, then decreasing the contents of chlorophyll (Wu et al., 2012). In line with this, Cd stress decreases the contents of chlorophyll (Fig.3D). When exposed to 50 μM CdCl_2_ or 75 μM CdCl_2_, in contrast to Col-0, the content of chlorophyll in *wrky45* plants is lower, and that in WRKY45-OX-15 plants is higher. The chlorophyll content in *wrky45* mutant is decreased by 7.8% and 13.1%, respectively, while it is increased by 7.5% and 28.3%, respectively, in WRKY45- OX-15 (P <0.05, Fig.3D). In short, these results confirmed that loss of *WRKY45* function leads to the decreased tolerance to Cd, while the overexpression of *WRKY45* enhances the Cd tolerance. This implies that *WRKY45* positively regulates Cd tolerance.

### 3.3 Effect of *WRKY45* on cadmium and iron accumulation

In order to test effects of *WRKY45* on Cd accumulation, we measured the Cd contents in roots and leaves of Col-0, *wrky45* and WRKY15-OX-15 plants under Cd stress. As to Cd contents in leaves, no difference can be found among Col-0, *wrky45* and WRKY45- OX-15 plants, whereas the contents of Cd in WRKY45-OX-15 roots are the highest, followed by Col-0 and *wrky45* plants (one way ANOVA, P < 0.05, Fig. 4A).

**Fig. 4.**
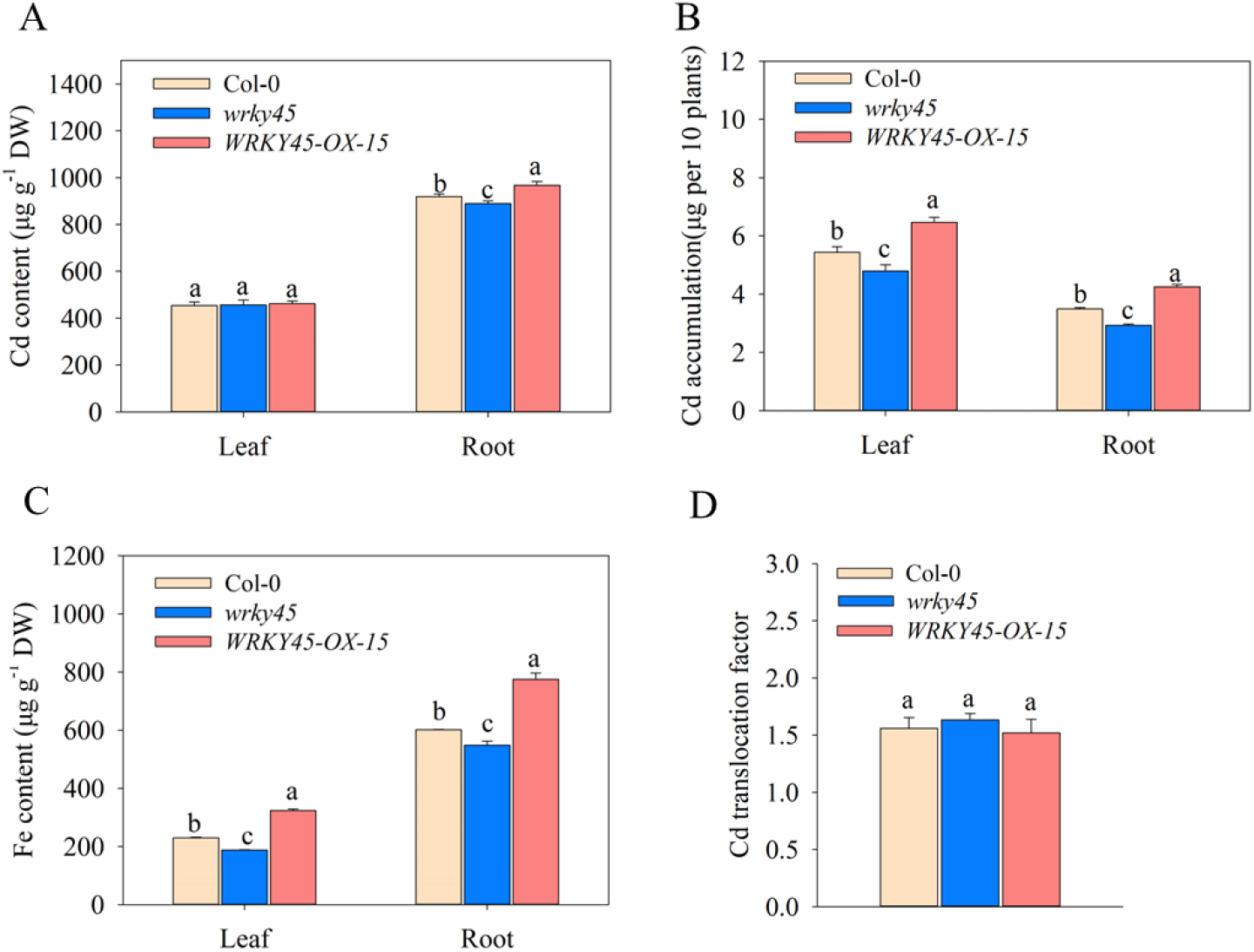
The Cd and Fe contents in leaves and roots of wild-type (Col-0), *wrky45* and overexpressing *WRKY45* (WRKY45-OX-15) lines treated with CdCl_2_. Seedlings were grown on hydroponic half strength MS nutrient solution for 2 weeks, and after treating them with 0 or 20 µM CdCl_2_ for 24 h, then the contents of Cd (A), Cd accumulation (B), and Fe (C) and Cd translocation factor (D) were measured. Results were means ± SE (n = 3). Different letters indicate significant difference at P < 0.05 level (one-way ANOVA with Turkey’s test).

Therefore, we calculated Cd accumulation (total Cd amount in plants). Compared with Col-0, the *wrky45* mutant exhibits reduced root and leaf Cd accumulation, with a decrease of 16.0% and 12.0%, respectively (P < 0.05). Conversely, the WRKY45-OX-15 plants display higher Cd accumulation than Col-0, with an increase of 21.9% in roots and 18.8% in leaves, respectively (P < 0.05, Fig. 4B). We defined the Cd translocation factor as the ratio of Cd accumulation in leaves to that in roots. We subsequently determine the Cd translocation factor under Cd stress and observed no difference among them (Fig. 4D). These results suggest that *WRKY45* regulates Cd tolerance in Arabidopsis not by reducing Cd uptake or translocation.

Given that the major root iron transporter *IRT1* also transports Cd, and considering the competition between Fe and Cd (Andres and Andrea, 2002; Clemens, 2006; Korshunova et al., 1999), We hypothesized that *WRKY45* affects the accumulation of iron ions and thus measured the iron content in WT, *wrky45*, and WRKY15-OX-15 plants under Cd stress. As shown in Fig. 4C, in contrast to Col-0, the contents of iron in *wrky45* leaves and roots are lower (P < 0.05), while higher in WRKY45-OX-15 leaves and roots (P < 0.05, Fig.4C). These findings indicated that *WRKY45* enhances iron uptake under Cd stress.

### 3.4 *WRKY45* positively regulates cadmium tolerance via glutathione-dependent PC synthesis pathway

The GSH-dependent pathway for PC synthesis has been documented as a crucial mechanism for heavy metal detoxification in plants (Cobbett et al., 1998; DalCorso et al., 2008; Kühnlenz et al., 2014; Mendoza-Cózatl et al., 2005). Therefore, we exploited buthionine sulfoximine (BSO), a glutathione synthesis inhibitor, to investigate whether *WRKY45* regulates Cd tolerance through the glutathione-dependent PC synthesis pathway. Col-0, *wrky45* and WRKY45-OX-15 seedlings exhibited comparable growth vigor when cultivated on 1/2 MS medium with 100 μM BSO (Fig. 5A). When treated with 50 μM Cd^2+^ plus 100 μM BSO or 75 μM Cd^2+^ plus 100 μM BSO, compared with Col-0, the *wrky45* mutant exhibited 9.9% or 57.1% reduction in primary root length, respectively (P < 0.05, Fig. 5B), 40.6% or 30.0% reduction in fresh weight (FW), respectively (P < 0.05, Fig. 5C), and 2.2% or 7.7% reduction in chlorophyll content, respectively (P < 0.05, Fig. 5D). When compared to Col-0, no significant differences were observed between WRKY45-OX-15 and Col-0 under the same conditions (Fig.5B- D). The results suggest that the involvement of *WRKY45* in Cd tolerance is dependent on endogenous GSH synthesis.

**Fig. 5.**
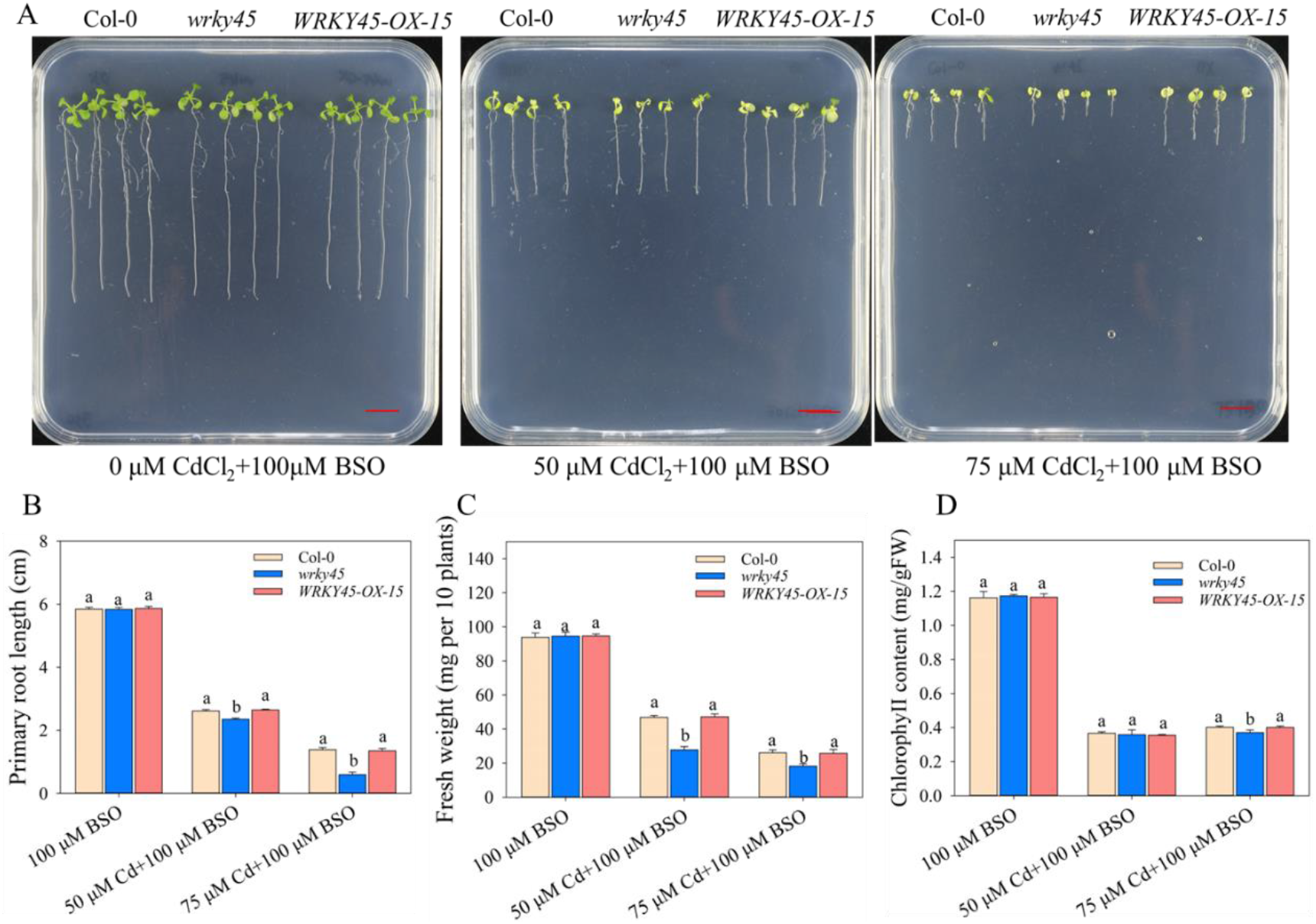
*WRKY45* regulates Cd tolerance through the GSH-dependent pathway. (A) Effect of GSH on growth of wild-type (Col-0), *wrky45* and WRKY45-OX-15 plants. Two-day-old seedlings grown on 1/2 MS media were transferred to 1/2 MS medium with or without 75μM CdCl_2_ or 150 μM GSH for 10 d. Scale bar, 1 cm. Effect of GSH on root length (B), fresh weight (C) and chlorophyll content (D) of Col-0, *wrky45* and WRKY45-OX-15 plants. Three independent experiments were carried out with similar results. Four seedlings per genotype from one plate were measured for each repeat. Data present the means ± SE (n = 4). Different letters indicate significant difference at P < 0.05 level (one-way ANOVA with Turkey’s test).

Furthermore, we treated with Arabidopsis seedlings with 75 μM Cd^2+^ and 150 μM GSH simultaneously. As to primary root length, in contrast to Cd application, addition of GSH reduces the toxicity of Cd in Col-0, *wrky45* and WRKY45-OX-15 seedlings, and the effects of GSH is most obvious in *wrky45* mutant (Fig.6A, 6B). As shown in Fig.6C, the application of GSH promotes the FW of *wrky45* plants (Fig.6C), even no difference between Col-0 and *wrky45* plants in 75 μM Cd^2+^plus 150 μM GSH conditions. With regards to chlorophyll contents, similar trend can be observed (Fig.6D). These findings indicate that the upregulation of *WRKY45* enhances Arabidopsis tolerance to Cd at least in part through a GSH-dependent approach. It seems to be that *wrky45* plants have lower endogenous levels of GSH and/or PC compared to Col-0 under Cd stress conditions.

**Fig. 6.**
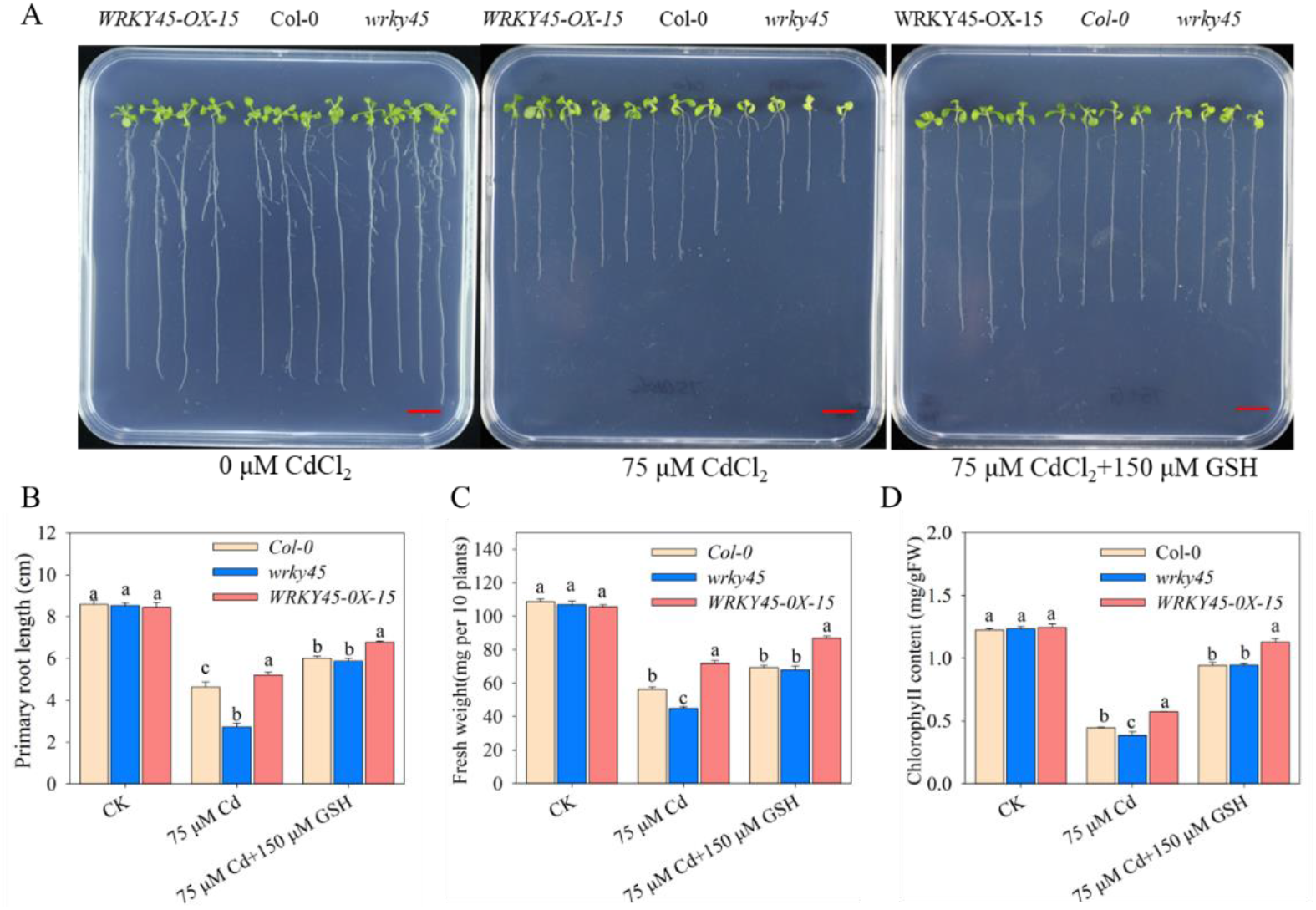
The Cd tolerance mediated by *WRKY45* is dependent on endogenous glutathione biosynthesis. (A) Effects of buthionine sulfoximine (BSO) on growth of Col-0), *wrky45* and WRKY45-OX lines. Seedlings were grown on half-strength Murashige and Skoog (1/2 MS) medium with 100 μM BSO, 100 μM BSO +50 µM Cd and 100 μM BSO +75 µM Cd for 10 days. Scale bar, 1 cm. (B-D) Effect of BSO on root length (B), fresh matter (C) and chlorophyll content (D) of Col-0, *wrky45* and WRKY45-OX plants described in A. Three independent experiments were done with similar results. Four plants per genotype from one plate were measured for each repeat. Data present the means ± SE (n = 4). Different letters indicate significant difference at P < 0.05 level (one-way ANOVA with Turkey’s test).

### 3.5 *WRKY45* promotes PCs production

It has been reported that exposure to Cd stress increases the levels of GSH and PCs in plants (Chen et al., 2015; Kühnlenz et al., 2014). Hence, we measured the levels of non-protein thiol peptides (NPT), GSH and phytochelatins (PCs) in Col-0, *wrky45*, and WRKY45-OX-15 plants.

No significant differences in NPT were observed among Col-0, *wrky45*, or WRKY45-OX-15 plants grown in Cd-free media (Fig. 7A). Under Cd stress, the NPT content is increased in Col-0, *wrky45* and WRKY45-OX-15 plants. The content of NPT in *wrky45* is decreased by 11.4% compared to that in Col-0 (P< 0.05), while the content in WRKY45-OX-15 plants is augmented by 7.1% (P < 0.05, Fig. 7A).

**Fig. 7.**
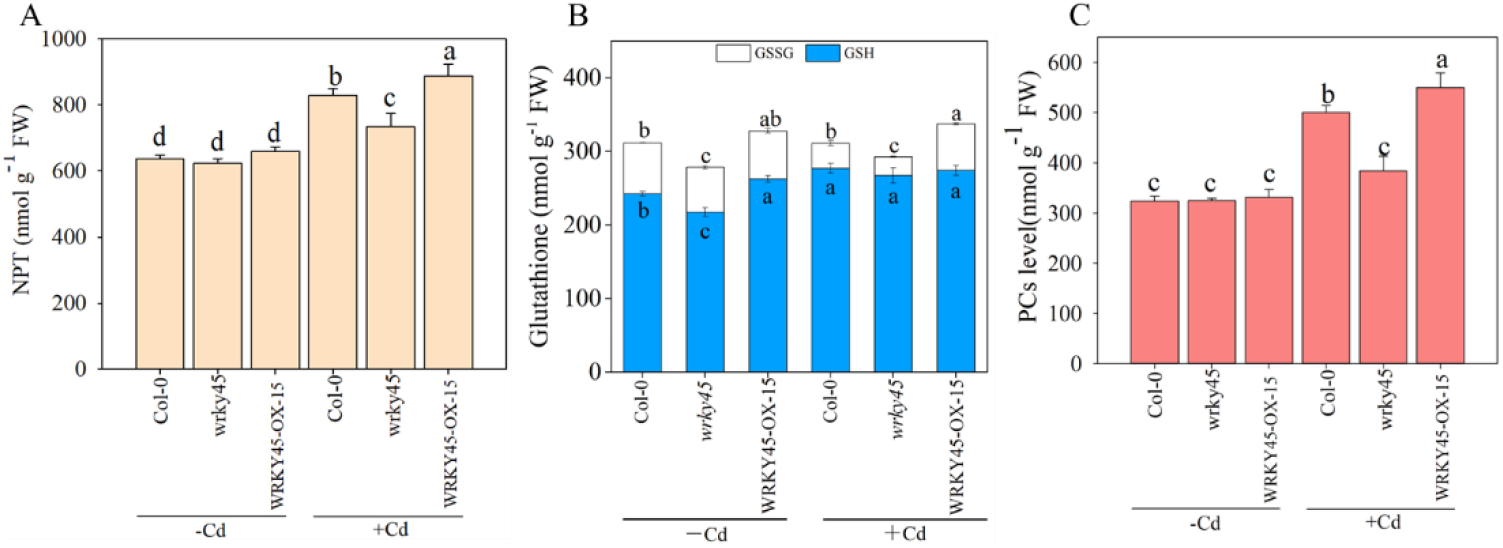
*WRKY45* promotes GSH and PCs production. (A-C) Measurements of non-protein thiol peptides (NPT)(A), total glutathione (GSH) plus 2 glutathione disulphide [GSSG] (B), and PC (C), contents in Col-0, *wrky45*, and the WRKY45-OX plants. Seedlings were grown on hydroponic nutrient solution for 2 weeks, and after treating them with 0 or 20 µM CdCl_2_ for 24 h, then the contents of NPT(A), GSH/GSSG (B), and PC (C) contents were quantified. Data present the means ± SE (n = 4). Different letters indicate significant difference at P < 0.05 level (one-way ANOVA with Turkey’s test). In Fig. 7.B. different letters in the upper section indicate significant difference in total glutathione, while letters in the lower section signify differences in reduced glutathione.

In addition, Cd stress induced an increase in the level of reduced GSH in plants, while having no impact on oxidized glutathione (GSSG) and total glutathione (GSH+GSSG). However, under both Cd-free or Cd stress conditions, WRKY45-OX-15 plants exhibit higher total glutathione content than Col-0, with 5.2% and 6.4% increase (P < 0.05, Fig. 7B). Conversely, *wrky45* mutants display a lower total glutathione content compared to Col-0, with a reduction ranging from 5.7% and 0.7% (P < 0.05, Fig. 7B).

There was no significant difference in PC levels among Col-0, *wrky45* and WRKY45- OX-15 plants under Cd-free conditions. However, under Cd stress conditions, the PC content is significantly increased in Col-0, *wrky45* and WRKY45-OX-15. Specifically, compared to Col-0, the PC content of *wrky45* plants decreased by 25.7% while increasing by 6.4% in WRKY45-OX-15 plants (P < 0.05, Fig. 7C). These results suggest that WRKY45 positively regulates PC production in Cd stress.

### 3.6 *WRKY45* promotes the expression of PC synthesis-related genes

Based on the above results (Fig. 6, Fig. 7), we speculated that transcription factor *WRKY45* appear to regulate the expression of PC synthesis-related genes. As revealed, Arabidopsis *GSH1*, *GSH2*, *PCS1* and *PCS2* are involved in PC biosynthesis, *GSH1* and *GSH2* generate the production of *GSH*, and *PCS1* and *PCS2* catalyze the production of PC (Han et al., 2019). We first investigated the responses of *GSH1*, *GSH2*, *PCS1*, and *PCS2* to Cd. As shown in Figure S5A and S5B, no significant differences were observed in the transcriptional levels of *GSH1* and *GSH2* among Col-0, *wrky45*, and WRKY45- OX-15 plants. Notably, the transcript levels of *PCS1* and *PCS2* are almost indistinguishable among Col-0, *wrky45*, and WRKY45OX-15 at 0 h (Cd treatment); however, after 6 h of Cd treatment, the levels of *PCS1* increase in all three genotypes with the highest abundance observed in WRKY45OX-15 plants followed by Col-0 and *wrky45*. (Fig.8A). Furthermore, a similar trend is observed in the expression of *PCS2*, with its transcription being induced by 6 hours of Cd treatment. Notably, the highest level of *PCS2* transcripts is detected in WRKY45 OX-15 plants, while the lowest level is found in *wrky45* mutant plants (Fig.8B). Our findings suggest that *WKKY45* promotes the upregulation of *PCS1* and *PCS2* in response to Cd stress, while exerting no influence on *GSH1* and *GSH2* expression.

**Fig. 8.**
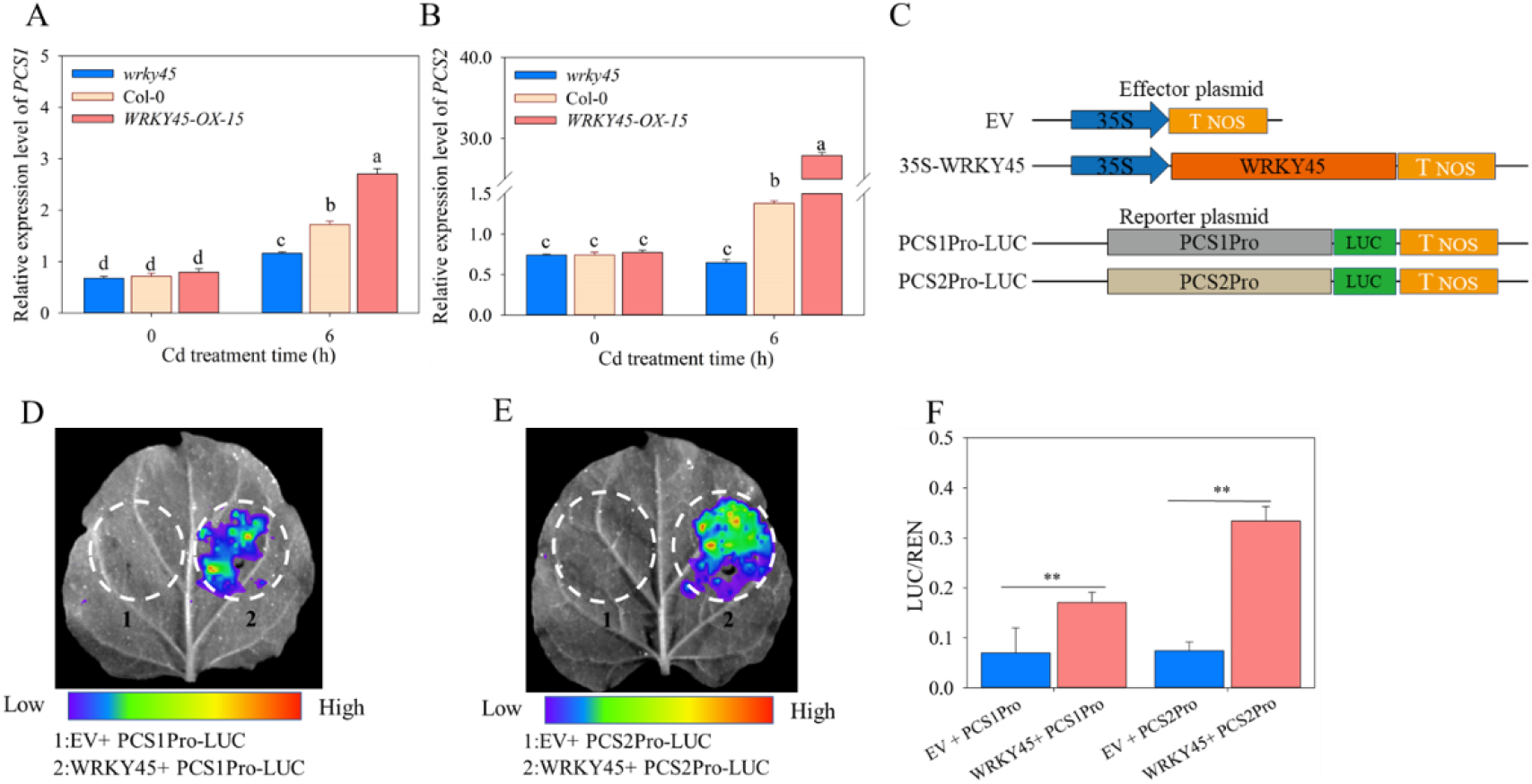
*WRKY45* promoted the expressions of *PCS1* and *PCS2* that catalyze the biosynthesis of phytochelatin (PCs) *in vivo*. The expressions of *PCS1* (A), and *PCS2* (B) in Col-0, *wrky45*, and WRKY45-OX lines by qRT-PCR. Two-week-old seedlings were treated with 20 µM CdCl_2_ for 0 h and 6 h. The relative expression was double normalized against *ACTIN2* (*AT3G18780*). (C) Schematics of all constructs used for transient expression assays in *Nicotiana benthamiana* leaves. The *PCS1* or *PCS2* promoter was fused to the luciferase (LUC) reporter gene, respectively. 35S:WRKY45 acted as an effector, and the empty vector (EV) acted as the negative control. (D, E) LUC fluorescence imaging showed expressions of PCS1Pro and PCS2Pro after co-expression with WRKY45. (F) The relative LUC/REN reporter activity was quantified. Data are means ± SE (n = 4). Different letters indicate significant difference at P < 0.05 (one-way ANOVA with Turkey’s test), and the asterisks indicate significant differences from the control (P < 0.05, Student’s *t*-test).

As reported, Arabidopsis WRKY regulate the expression of downstream genes via binding the W-box *cis*-elements (Rushton et al., 2010). Hence, we screened the W-box elements in the promoter regions of *PCS1* and *PCS2* and confirmed the existence (Fig.9A). We thus deduced that *WRKY45* plays a direct role in regulating the transcriptional activity of both *PCS1* and *PCS2* in Arabidopsis. To verify this, we cloned the native promoter of *PCS1* and *PCS2*, respectively, and fused them with a luciferase reporter gene (Fig. 8C). We then co-transformed *Nicotiana benthamiana* with the plasmid expressing *WRKY45* driven by 35S promoter. Fluorescence imaging showed that WRKY45 promotes the expression of both PCS1Pro-LUC and PCS2Pro-LUC (Fig. 8D, E), and the expression of *WRKY45* increased the LUC/REN ratio to much higher extent than the empty vector in *PCS1* and *PCS2* promoter with different effect (P < 0.01), the promotive effects on PCS2 promoter is stronger than PCS1 promoter (Fig. 8F).

**Fig. 9.**
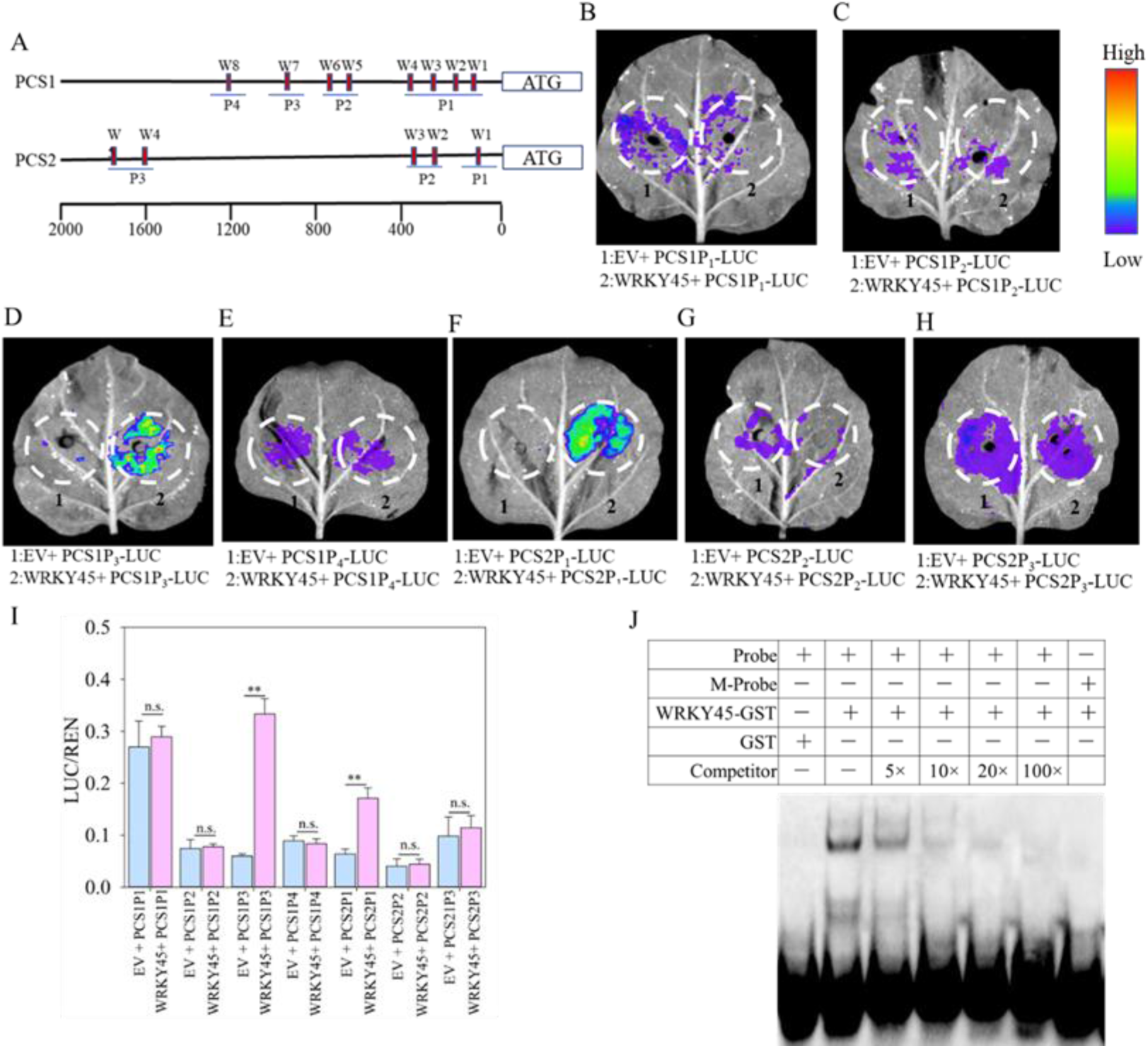
*WRKY45* promoted the expression of *PCS1* and *PCS2* by directly binding its promoter *in vivo*. (A) WRKY45-binding site (W-BOX, TTGAC[C/T]) prediction within 2,000 bp of *PCS1* and *PCS2* promoters. The PCS1 promoter was divided into four segments, namely PCS1 probe 1 (PCS1-P1), PCS1-P2, PCS1-P3 and PCS1-P4. The PCS2 promoter was divided into three segments, namely PCS2 probe 1 (PCS2-P1), PCS2-P2 and PCS2- P3. (B-H) LUC fluorescence imaging showed that WRKY45 activated PCS1-P3 (D) and PCS2-P1 (F), respectively. WRKY45 did not bind PCS1-P1, PCS1-P2, and PCS1-P4 (B-E), PCS2-P2 (G) and PCS2-P3 (H). (I) The relative LUC/REN reporter activity was quantified. PCS1P1-LUC, PCSP2-LUC, PCS1P3-LUC, PCS1P4-LUC, PCS2P1-LUC, PCS2P2-LUC, or PCS2P3-LUC was co-transformed with 35S:WRKY45 into *N.benthamiana*. (J) Electrophoretic mobility shift assay (EMSA) showed that WRKY45 protein bounds to W-box1(W1) in PCS2-P1 probe, but not mutated W-box1 (mW-box1). Data are presented as means ± SE (n = 4). The asterisks indicate significant differences from the control (P < 0.05, Student’s *t*-test).

The binding of WRKY to the core W-box (TTGACC/T) motif in the promoter region of downstream target genes is a well-established fact (Ciolkowski et al., 2008). The promoter regions of *PCS1* and *PCS2* were found to contain W-box *cis*-elements. (Fig. 9A). We postulated that WRKY45 binds directly to these W-boxes and exerts its impact on their expression. To test the hypothesis, four fragments of native PCS1 promoter region were cloned, each containing W-box and designated as PCS1P1 (-90— 276 bp), PCS1P2 (-406—715 bp), PCS1P3 (-685—815 bp), and PCS1P4 (-1249—1431 bp) upstream of the initiation codon ATG (Fig. 9A). These fragments were then individually fused to a luciferase reporter gene and transfected into *Nicotiana benthamiana* along with *WRKY45* driven by 35S promoter. Fluorescence imaging revealed that *WRKY45* specifically enhances luciferase reporter gene expression driven by the P3 fragment of *PCS1* promoter, while has effects on P1, P2 or P4 (Fig. 9D). Consistently, the overexpression of *WRKY45* significantly enhanced luciferase activity compared to the vector control in PCS1P3 (Fig. 9I).

The 2200 bp-long native promoter region of *PCS2*, which contains W-box, can be divided into three segments: PCS2P1 (-226-0bp), PCS2P2 (-486--210 bp), and PCS2P3 (- 1870--2108bp). Using the same method, we have also demonstrated that *WRKY45* enhances the expression of a reporter gene under the control of P1 segment from *PCS2* promoter (Figure 9F).

Furthermore, our EMSA assay verified the direct binding of WRKY45 protein to the biotin-labeled W-box1 probe located in the P1 segment of *PCS2* promoter, with almost complete inhibition upon addition of 20-fold excess unlabeled competitive no-labelled cold probes (Fig. 9J). In addition, WRKY45 fails to bind to the mutated W-box probe mW-box ("CGCGGA"). These EMSA results confirmed that *WRKY45* regulates the transcription of *PCS2* by directly binding to the W-box in the *PCS2* promoter region *in vitro*. On the other hand, we tried to confirm the interaction between *WRKY45* and the W-box motif in *PCS1* via EMSA analysis; but failed. The possible reason might be that the flanking sequence of W-box play roles in the binding of WRKY45 with *PCS1* W-box element.

### 3.7 *WRKY45* genetically acts upstream of *PCS1* and *PCS2* to positively regulate Cd tolerance

As we provided biochemical evidence to verify the interaction between *WRKY45* and *PCS1* or *PCS2* (Fig.9). *PCS1* and *PCS2* are the direct targets of *WRKY45*, which confers *Arabidopsis tolerance* by modulating their expression. We tried to provide genetic evidences. *PCS1* or *PCS2*-related mutants are more sensitive to Cd toxicity (Kühnlenz et al., 2014). We postulated that the *wrky45* mutant phenotype to Cd stress could be rescued through the overexpression of *PCS1* and *PCS2*, respectively. To this end, we introduced PCS1-OX and PCS2-OX into *wrky45* mutant. We screened the overexpressing *PCS1* or *PCS2* lines in *wrky45* background via qRT-PCR (Fig. 10A, Fig. S4A). Then the *wrky45*/PCS1-OX-1, *wrky45*/PCS1-OX-12, *wrky45*/PCS2-OX-6 and *wrky45*/PCS2-OX-11 plants were used further, respectively. In Cd-free half-strength MS media, the growth of *wrky45*/PCS1-OX-1, *wrky45*/PCS1-OX-12, *wrky45*/PCS2-OX-6, and *wrky45*/PCS2-OX-11 plants does not exhibit significant difference compared to Col-0 and *wrky45* mutant plants. However, based on primary root length (Fig. 10C, Fig. S4C), fresh weight (Fig. 10D, Fig. S4D) and chlorophyll content (Fig. 10E, Fig. S4E) in 1/2 MS media containing 75 μM Cd^2+^, the wrky*45* PCS1-OX and *wrky45* PCS2-OX plants are less sensitive to Cd compared to the *wrky45* mutants and even more tolerant to Cd than Col-0 plants. Collectively, these findings suggest that *WRKY45* functions upstream of *PCS1* and *PCS2* in the Cd response signaling pathway genetically.

**Fig. 10.**
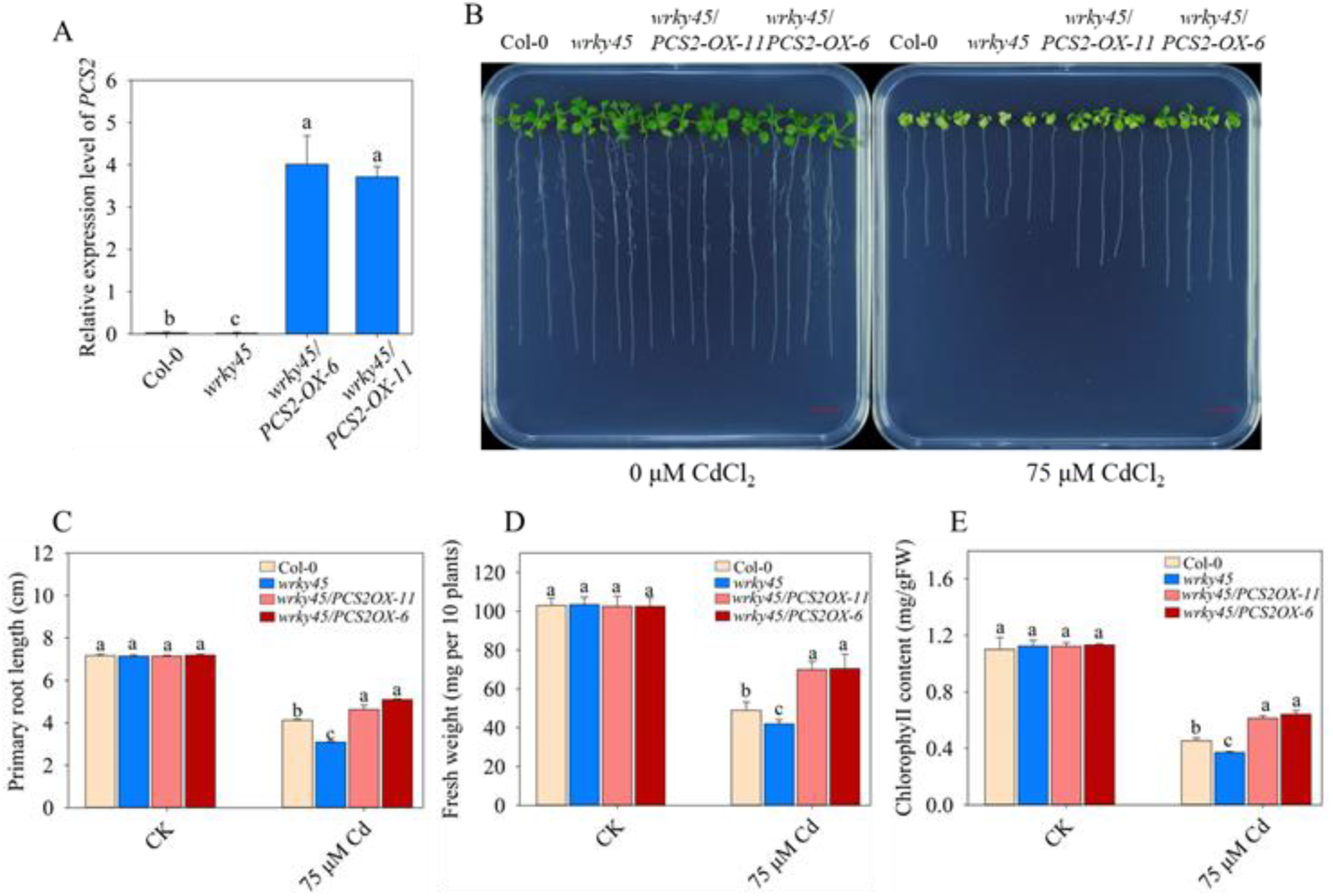
Overexpression of *PCS2* rescues the Cd-sensitive phenotypes of *wrky45* mutant. (A) Expression of *PCS2* of *wrky45*, wild-type (Col-0), *wrky45/*PCS2-OX-6, and *wrky45/*PCS1-OX-11 lines. (B) Growth of Col-0, *wrky45, wrky45/*PCS2-OX-6, and *wrky45/*PCS2-OX-11 lines under Cd stress. Two-day-old plants grown on half-strength Murashige and Skoog’s (1/2 MS) media were transferred to 1/2 MS media without or with 75 μM CdCl_2_. Photographs were taken 10 days after the transfer. Scale Bar, 1 cm. (C-E) Primary root length (C), fresh weight (D) and chlorophyll (E) of plants were measured. Three independent experiments were done with similar results, each with three biologicals repeats. Four plants per genotype from one plate were measured for each repeat. Data are presented as means ± SE (n = 4). Different letters indicate significant difference at P < 0.05 level (one-way ANOVA with Turkey’s test).

## 4. Discussion

Arabidopsis *WRKY45* transcription factors have been shown to be involved in the regulation of phosphorus uptake (Wang et al., 2014), leaf senescence (Chen et al., 2017; Barros et al., 2022). Here, we uncovered a novel function of *WRKY45* in conferring tolerance to Cd stress. We verified that the WRKY45-PCS module is required for Cd detoxification in *Arabidopsis thaliana.* We found that Cd stress significantly induces the expression of *WRKY45* and WRKY45 protein level (Fig.1), in line with this, overexpressing *WRKY45* line is more tolerant to Cd, while *wrky45* mutant is more sensitive to Cd (Fig. 3). *WRKY45* directly activated the transcription of *PCS1* and *PCS2* (Fig. 8; Fig.9), thus increasing PC synthesis (Fig. 7).

Transcription factor WRKYs have been reported to control plant development, pathogen defense and stress responses (Dong et al., 2003; Ulker and Somssich, 2004). Previous studies have shown that *WRKY12* (Han et al., 2019)*, WRKY13* (Sheng et al., 2019)*, WRKY33* plays roles in the response of plants to Cd toxicity (Zhang et al., 2020). Interestingly, the amino acid sequences of *WRKY45* and *WRKY33* are very similar, and they belong to the same subgroup in the phylogenetic tree of *Arabidopsis thaliana* WRKY (Eulgem et al., 2000). It was found that both *WRKY45* and *WRKY33* played a positive regulatory role in phosphorus nutrients (Shen et al., 2021; Wang et al., 2014). In addition, *WRKY45, WRKY33, WRKY13, WRKY12* also plays a certain role in regulating plant growth and development (Chen et al., 2017; Li et al., 2016; Sun et al., 2020). Like *WRKY45, WRKY75* also regulate leaf senescence and low phosphorus response (Devaiah et al., 2007; Zhang et al., 2018; Zhang et al., 2022). Considering *WRKY75* is the analogue of *WRKY45* in Arabidopsis, it is interesting to test the involvement of *WRKY75* in the adaptations to Cd stress.

WRKY proteins from various plant species such as *Arabidopsis thaliana*, *Glycine max*, *Zea mays L*., Populus, potato, wheat, tomato, *Thlaspi caerulescens* and *Tamarix hispida* are involved in Cd tolerance (Hong et al., 2017; Khan et al., 2023; Li et al., 2022; Liu et al., 2023; Wei et al., 2008; Wu et al., 2022; Wu et al., 2023; Xian et al., 2023). For example, *AtWRKY12* inhibits *GSH* expression via bind the W-box element of *GSH1* promoter, thus attenuating the tolerance to Cd (Han et al., 2019). In contrary, *AtWRKY13* positively regulates *PDR8* expression via directly binding the promoter of *PDR8* than encodes ABCG-type transporter, consistently overexpressing *WRKY13* decrease Cd accumulation and enhanced Cd tolerance (Sheng et al., 2019). *GmWRKY142* maintains Cd tolerance by up-regulating the Cadmium Tolerance 1-Like genes (Cai et al., 2020). In addition, the overexpression of *GmWRKY172* enhances the tolerance of soybean to Cd and reduces the contents of Cd in seeds (Xian et al., 2023). *ZmWRKY4* appear to govern the expression of *ZmSOD4* and *ZmcAPX* under Cd stress (Hong et al., 2017). The V-ATPase c subunit gene (*ThVHAc1*) in *Tamarix hispida* is the target of ThWRKY7 that bind to the W-box *cis*-element (Yang et al., 2016). The Populus *PyWRKY48* increases Cd tolerance via up-regulating *PaGRP, PaPER* and *PaPHOS*, which encode cell wall proteins, antioxidant enzyme, and heavy metal-associated proteins, respectively (Wu et al., 2023). Interestingly overexpression of *PyWRKY75* increases the Cd tolerance, Cd accumulation, and the contents of chlorophyll (Wu et al., 2022). Transcriptome analysis indicate some *Solanum lycopersicum* L. WRKY genes respond to Cd stress (Khan et al., 2023). Collectively, these results suggested that WRKY-modulated Cd resistance is conserved across plant species.

As shown in Figure 1, the transcripts of *WRKY45* are rapidly induced after 6 hours of Cd treatment (Fig.1). Therefore, it is intriguing to investigate which transcription factors regulates the expression of *WRKY45* in future studies. Additionally, the protein level of WRKY45 also increases upon Cd application (Fig.1C). Given that the possibility of post-translational modifications on *WRKY45*, it is imperative to explore the kinase and E3 ligase responsible for its phosphorylation and ubiquitination under Cd stress via Y2H and BiFC assays. In rice, the kinase responsible for phosphorylating *WRKY45* and the E3 ligase that facilitates turnover of *WRKY45* have been extensively investigated (Ichimaru et al., 2022).

Under Cd exposure, young leaves exhibit chlorosis, root growth is inhibited and eventually ceases (Kahle, 1993). As documented, the transition zone (TZ) and elongation zone (EZ) of primary root are the target site of Cd, the cell death caused by Cd stress taken place in TZ and EZ, moreover the growth inhibition of Cd in primary root is partially dependent on ethylene signaling (Kong *et al*., 2018). The disruption of chloroplast ultrastructure by Cd impedes the accumulation of photosynthetic pigments and reduces the rate of photosynthesis, ultimately compromising crop yield, quality, and fresh weight (López-Millán et al., 2009). In line with this, Cd stress inhibits primary root growth, decreases fresh weight and chlorophyll contents (Fig. 3). Notably, the mutation of *WRKY45* gene boosts Arabidopsis sensitivity to Cd stress, and significantly reduces primary root length, fresh weight matter, and chlorophyll content. Conversely, the overexpression of *WRKY45* effectively enhances Cd tolerance and significantly increases primary root length, fresh weight, and chlorophyll content in contrast to Col-0 plants (Fig. 3). Hence, it is imperative to identify the downstream genes of *WRKY45* that are involved in primary root growth, chlorophyll biosynthesis and degradation.

Reducing Cd accumulation through the inhibition of Cd absorption or promotion of Cd excretion is a crucial mechanism for plants to maintain Cd tolerance (Kim et al., 2007; Mendoza-Cozatl et al., 2011). Therefore, we conducted an analysis of Cd accumulation levels in the roots and leaves of Col-0, *wrky45* and *WRKY45-OX* line plants under Cd stress. Our findings indicate that overexpression of *WRKY45* significantly increases Cd content in Arabidopsis roots and leaves under Cd stress, while the opposite effect is observed in *wrky45* mutant plants (Fig. 4A). There was no significant difference observed in the Cd content of leaves, suggesting that the reduction of Cd uptake pathway is not a mechanism underlying WRKY45-mediated Cd tolerance. Moreover, owing to the chemical resemblance among heavy metals, Cd disrupts the dynamic equilibrium of other metal ions like iron (Nazar et al., 2012). Therefore, typical symptoms of iron deficiency were observed in plants subjected to Cd stress (Meng et al., 2022). The enhancement of iron nutrition facilitates the augmentation of plants’ tolerance and accumulation towards Cd (Zhu et al., 2020). In this study, we observed that the overexpression of *WRKY45* under Cd treatment significantly enhanced the Fe content in Arabidopsis roots and leaves, whereas a decrease in Fe content was detected in *wrky45* mutant plants in roots and leaves, respectively (Fig. 4C). Considering the competition of Cd and iron during uptake, and the important role of iron in chlorophyll biosynthesis, hence increasing iron acquisition and translocation of iron to leaves might be one of strategy of Arabidopsis to cope with Cd stress via *WRKY45*. Importantly, different from *AtWRKY13* that decrease the uptake of Cd in Arabidopsis, *WRKY45* does not decrease Cd uptake and translocation, but increase PC production and iron uptake and translocation.

The biosynthesis pathway of GSH-dependent plant PC synthesis is an important mechanism for promoting heavy metal tolerance through the chelation of GSH or PC with metals (Cobbett et al., 1998; DalCorso et al., 2008; Kühnlenz et al., 2014; Mendoza-Cózatl et al., 2005). For example, under Cd stress, plants chelate free Cd^2+^ in cells by synthesizing PC and isolate PC-Cd^2+^ in plant vacuoles to relieve Cd poisoning (Agarwal et al., 2020, Riaz et al., 2021; Tang et al., 2023; Zhao et al., 2022). We observed that the tolerance of WRKY45-OX-15 plants to Cd was abolished in half-strength media supplemented with 100 μM BSO plus 50 or 75 μM Cd^2+^, whereas *wrky45* mutant plants exhibit increased sensitivity to Cd (Fig. 5). As reported, application of BSO inhibits endogenous GSH biosynthesis (Kim et al., 2007). Hence it seems to be that BSO application decreased GSH content in overexpressing *WRKY45* line, thus resulting in the low level of PCs in *Arabidopsis.* Additionally, we found that the Cd sensitivity of *wrky45* mutant was alleviated in half-strength media supplemented with 150 μM GSH and 75 μM Cd^2+^, and the Cd tolerance of WRKY45-OX-15 plants was further enhanced (Fig. 6). Therefore, we speculate that WRKY45 may be involved in plant tolerance to Cd stress through GSH-dependent PC synthesis pathway. Further, we analyzed the PC levels of Col-0, *wrky45* and *WRKY45-OX* lines under Cd stress, and found that overexpression of *WRKY45* significantly increases the content of PC under Cd stress, while PC content of *wrky45* mutant decreased significantly (Fig. 7C). These results indicates that *WRKY45* regulated Cd tolerance in Arabidopsis via GSH-PCs synthesis pathway.

PC is a low molecular weight peptide thiol group synthesized by GSH via PC synthase (PCS) (Cobbett and Goldsbrough., 2002). The synthesis of PC in *Arabidopsis thaliana* is predominantly mediated by PCS1 and PCS2 (Howden et al., 1995; Kühnlenz et al., 2014). PCS1is the major form of PCS in Arabidopsis (Howden et al., 1995). We ordered and identified the *cad1-3* mutant of *AtPCS1* and the *pcs2* mutant of *AtPCS2* (Fig. S2). Consistent with previous studies (Kühnlenz et al., 2014), *cad1-3* and *pcs2* are more sensitive to Cd toxicity (Fig. S3). In *Arabidopsis thaliana*, the transcription factor *ZAT6* directly upregulates the transcription of *GSH1*, and MYB4 promotes the transcription of *PCS1* (Agarwal et al., 2020; Chen et al., 2016). *WRKY12* binds to the W-box element in the promoter of *GSH1*, thereby repressing its transcriptional activity, additionally, it indirectly downregulates other genes involved in PC synthesis (*GSH2, PCS1* and *PCS2*), thus plays negative roles in Cd tolerance in Arabidopsis (Han et al., 2019). In this study, we identified the presence of W-box elements within the 2000 bp promoter region upstream of the start codon of *PCS1*, and *PCS2*, respectively (Fig. 9A).

qRT-PCR analysis revealed that *WRKY45* has positive regulatory effect on the transcription of *PCS1* and *PCS2* (Fig. 8AB), as the transcripts of *PCS1* and *PCS2* in WRKY45-OX-15 plants are more than that in *wrky45* mutant. Dual-Luc experiment analysis further confirmed that *WRKY45* is the transcription activator of *PCS1* and *PCS2* (Fig. 8F). In order to elucidate the precise binding sites of *WRKY45* on *PCS1* and *PCS2* promoters, we cloned these promoters containing W-box regions into four (P1, P2, P3 and P4) and three (P1, P2, and P3) segments respectively (Fig. 9A). Dual fluorescence assay revealed that WRKY45 activates PCS1P3 and PCS2P1 regions with high specificity (Fig.9I). We synthesized 28-bp-long probes containing W-box7 (TTGACT) in PCS1P3 and W-box1 (TTGACT) in PCS2P1, which were subsequently labeled with biotin. EMSA analysis revealed that WRKY45 binds to the W-box1 region of *PCS2* promoter (Fig. 9J), but not *PCS1*. Despite we repeated EMSA experiments, we were unable to detect the interaction between WRKY45 and the *PCS1* W-box7 element. This may be attributed to the flanking sequence of the W-box *cis*-element, which appears to affect binding with WRKY45.

Genetic analysis showed that the overexpression of *PCS1* and *PCS2* in *wrky45* increase the tolerance of mutant to Cd toxicity, indicating that *PCS1* and *PCS2* act downstream of *WRKY45* (Fig. 10, Fig. S4). Although we have demonstrated that WRKY45 positively regulates GSH synthesis in plants (Fig. 7B), *WRKY45* has no transcriptional regulation on GSH synthase genes *GSH1* and *GSH2*, respectively (Fig. S5). It seems to be possible that *WRKY45* may indirectly influence the Cd-mediated GSH synthesis via the upstream genes. Collectively, our findings indicate that *WRKY45* modulates Arabidopsis Cd tolerance via a GSH-dependent PC synthesis pathway.

The induction of both *WRKY45* expression and protein levels under Cd conditions highlights the importance of identifying upstream transcription factors that directly regulate *WRKY45* transcription, as well as proteins involved in its degradation. In addition, further research should be conducted on the function of *WRKY45* in other heavy metals such as iron, copper, mercury, and lead stress. More importantly, deciphering the pathway and networks mediated by WRKY in crops in Cd-contaminated land is urgent in agriculture.

## Conclusion

Apart from the functions of Arabidopsis WRKY45 in phosphorus nutrition and leaf senescence, we herein discovered a novel function of *WRKY45* in response to Cd stress. We proposed a working model for the roles of *WRKY45* in conferring Cd tolerance. The transcription factor *WRKY45* induced quickly by Cd stress directly activates the expression of genes encoding PC biosynthesis enzymes, *PCS1* and *PCS2*, thus increasing PC content and enhancing Cd tolerance. In addition, the overexpression of *WRKY45* promotes iron uptake under Cd stressful conditions (Fig. 11). Taken together, we provided physiological, biochemical, and genetic evidences to verify that Arabidopsis WRKY45 confers Cd tolerance in *Arabidopsis* via activating *PCS1* and *PCS2* expression.

**Fig. 11.**
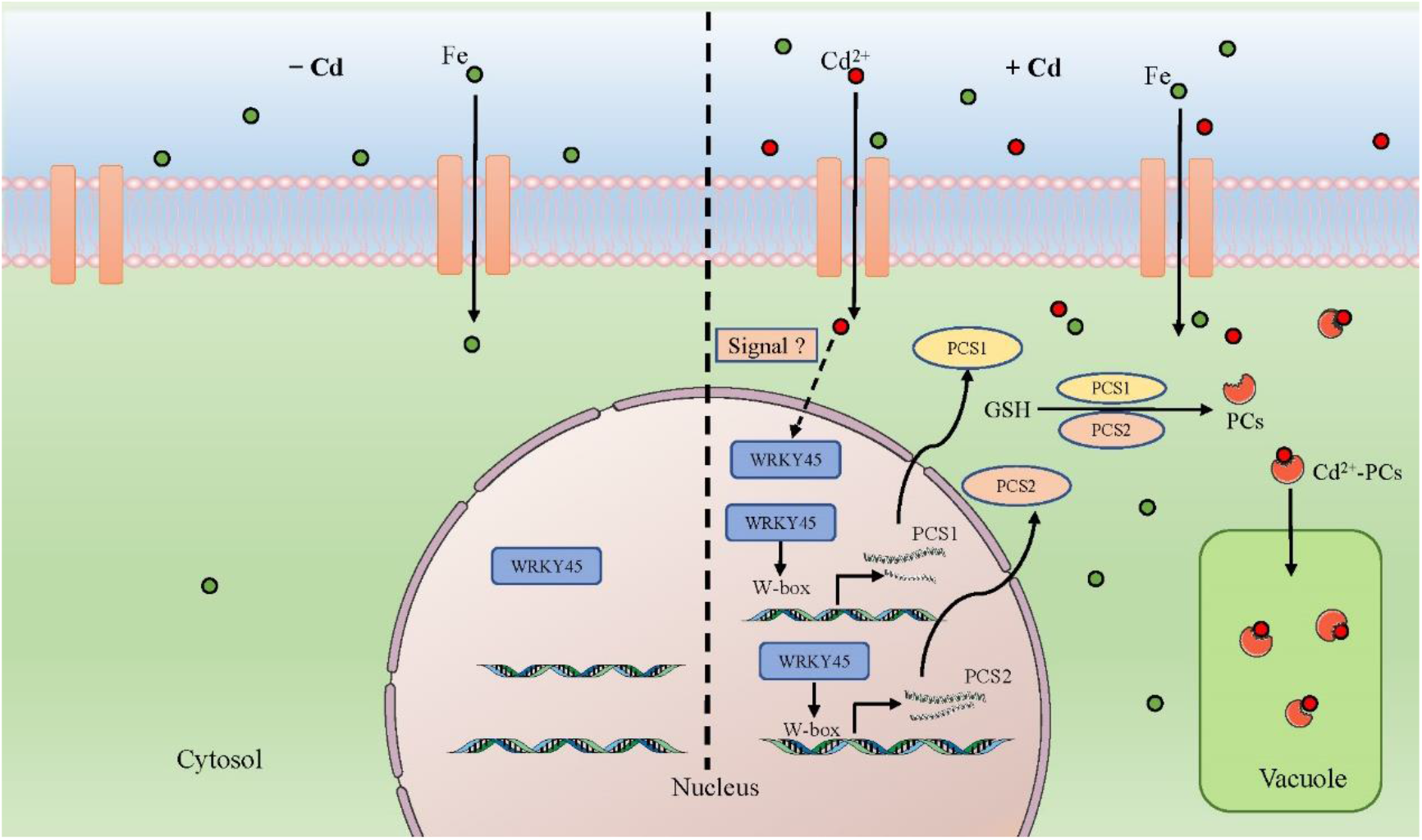
The working model about the roles of *WRKY45* in regulating Cd tolerance. Cd stress promotes the levels of *WRKY45*, which directly activates the transcription of plant *phytochelatin synthase 1* (*PCS1*) and *PCS2*, thus increasing the accumulation of plant phytochelatin. In addition, overexpressing increase the uptake of iron from media. Hence, WRKY45 confers Cd tolerance in Arabidopsis. Red ball and green ball indicate Cd and Fe, respectively.

### CRediT authorship

Fangjian Li: Conceptualization, Methodology, Validation, Formal analysis, Investigation, writing original draft, Writing review & editing. Yaru Deng: Validation. Yan Liu: Validation. Cuishan Mai: Validation. Yun Xu: Formal analysis. Jiarui Wu: Formal analysis. Cuiyue Liang: Review. Jinxiang Wang: Supervision, Conceptualization, Funding acquisition, Project administration, Resources, Writing – review & editing.

## Data Availability

Data will be made available on request.

## Acknowledgements

This study was partially supported by the Science and Technology Planning Project of Guangdong Province of China Foundation (Grant No. 2021B1212040008). We thank Dr. Diqiu Yu (Chinese Academy of Sciences, China) and ABRC (https://abrc.osu.edu/)) for providing Arabidopsis seeds, and Dr. Jiang Tian (South China Agricultural University) for help and discussions.

---

**Fig. S1.**
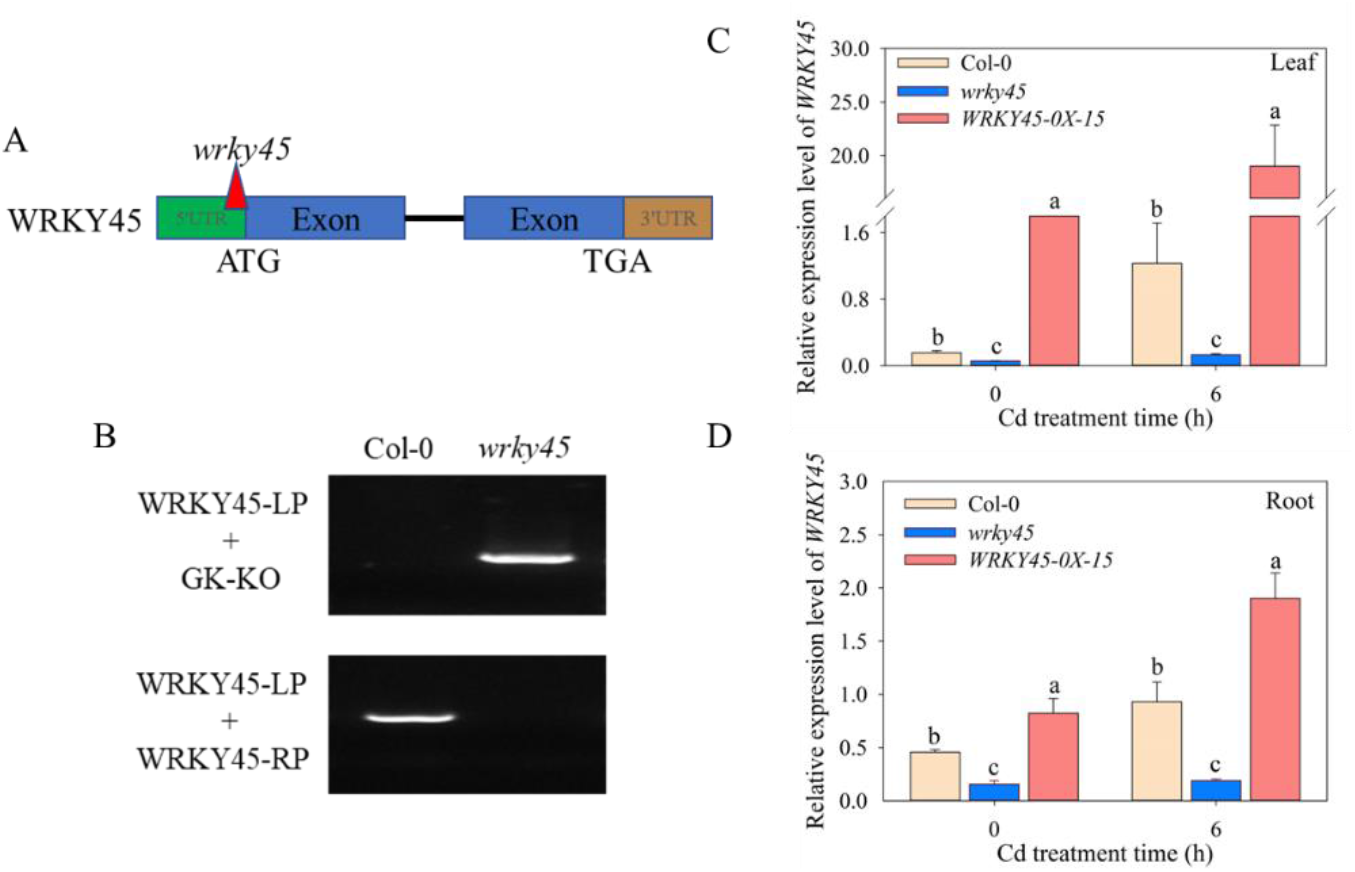
Verification of *WRKY45* knock-down in *wrky45* seedlings; (A) Position of T-DNA insertion in the wrky45 mutant. (B) Genomic PCR genotyping of the DNA of the *wrky45* mutant. Primers used for genomic PCR are listed in Supplementary table S1. (C, D) Expression of *WRKY45* in the leaves and roots of wild-type (Col-0), *wrky45* and WRKY45-OX-15 plants. qRT-PCR primers are listed in Supplementary table S1. Seedlings were grown on hydroponic nutrient solution for 2 weeks, and after treating them with 20 µM CdCl_2_ for and then analyzed at 0 h, 6 h after seedling transfer. Data are means ± SE (n = 4). Different letters indicate significant difference at P < 0.05 level (one-way ANOVA with Turkey’s test).

**Fig. S2.**
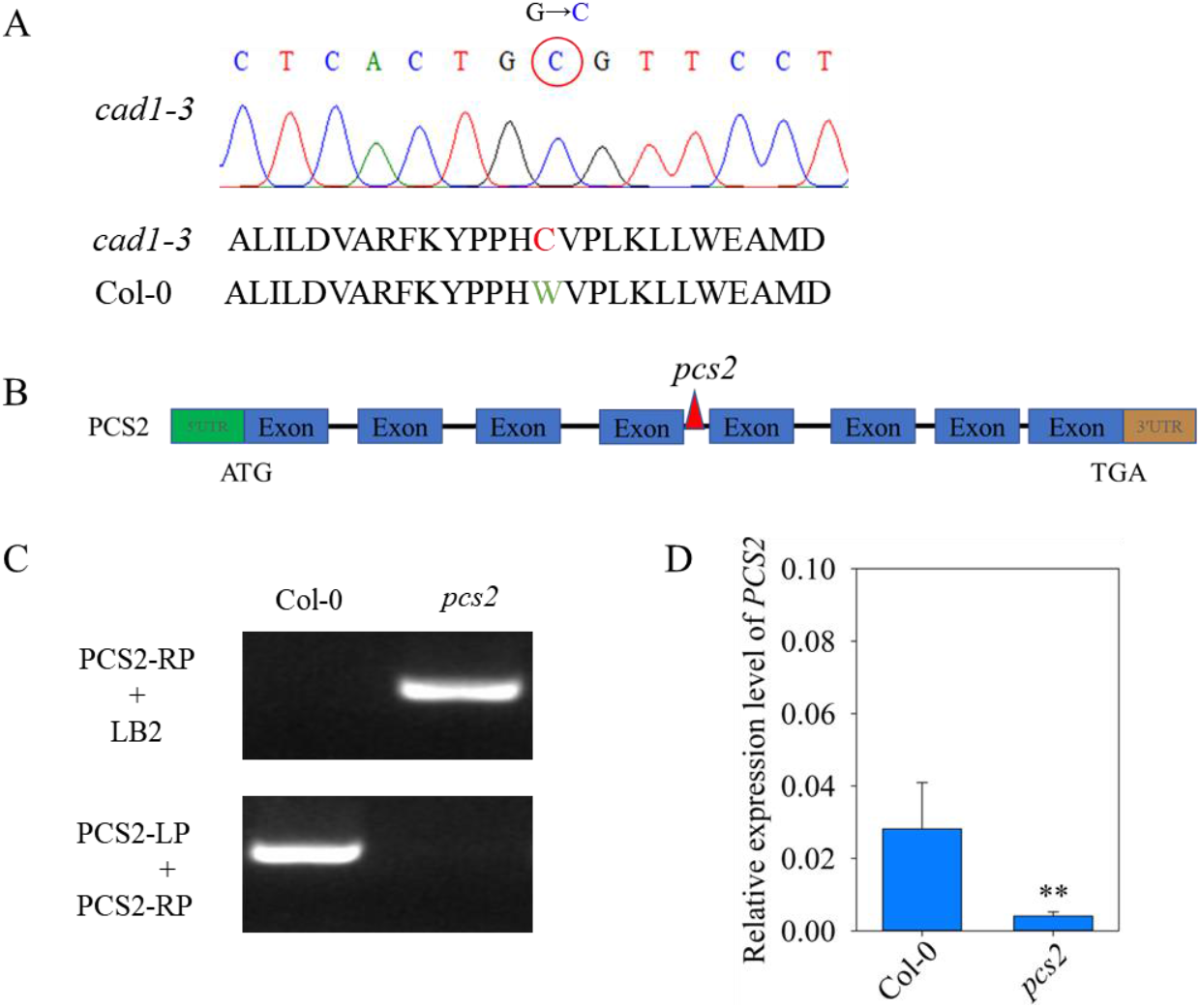
Identification of *cad1-3* and *pcs2* mutants; (A) Sequencing results of *cad1-3* mutant. (B) Position of T-DNA insertion in the *pcs2* mutant. (C) Genomic PCR genotyping of the DNA of the *pcs2* mutant. Primers used for genomic PCR are listed in Supplementary table S1. (D)Expression of *PCS2* in the wild-type (Col-0), and *pcs2* mutants. qRT-PCR primers are listed in Supplementary table S1. Bars in the graph show mean ± SE calculated from 4 biological replicates. Data are means ± SE (n = 4). The asterisks indicate significant differences from the control (P < 0.05, Student’s *t*-test).

**Fig. S3.**
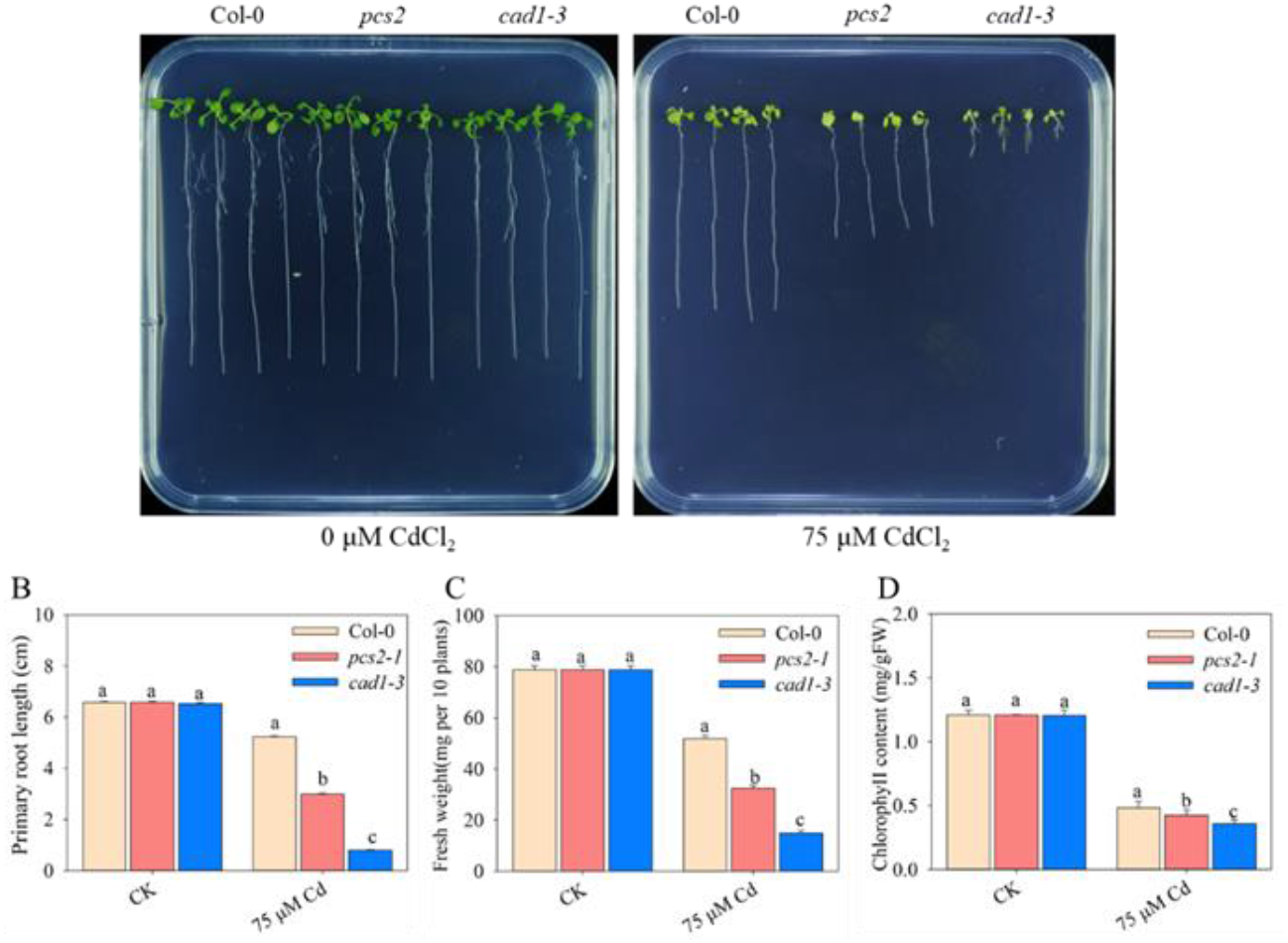
The *cad1-3* and *pcs2* mutant plants are more sensitive to cadmium. (A) Growth of Col-0), *cad1-3*, and *pcs2* plants under Cd stress. Two-day-old plants grown on half-strength Murashige and Skoog’s (1/2 MS) medium were transferred to 1/2 MS medium without or with 75 μM CdCl_2_. Photographs were taken 10 days after the transfer. Scale bar, 1 cm. (B-D) Primary root length (B), fresh weight (C) and chlorophyll (D) were measured. Three independent experiments were done with similar results, four plants per genotype from one plate were measured for each repeat. Data are means ± SE (n = 4). Different letters indicate significant difference at P < 0.05 level (one-way ANOVA with Turkey’s test).

**Fig. S4.**
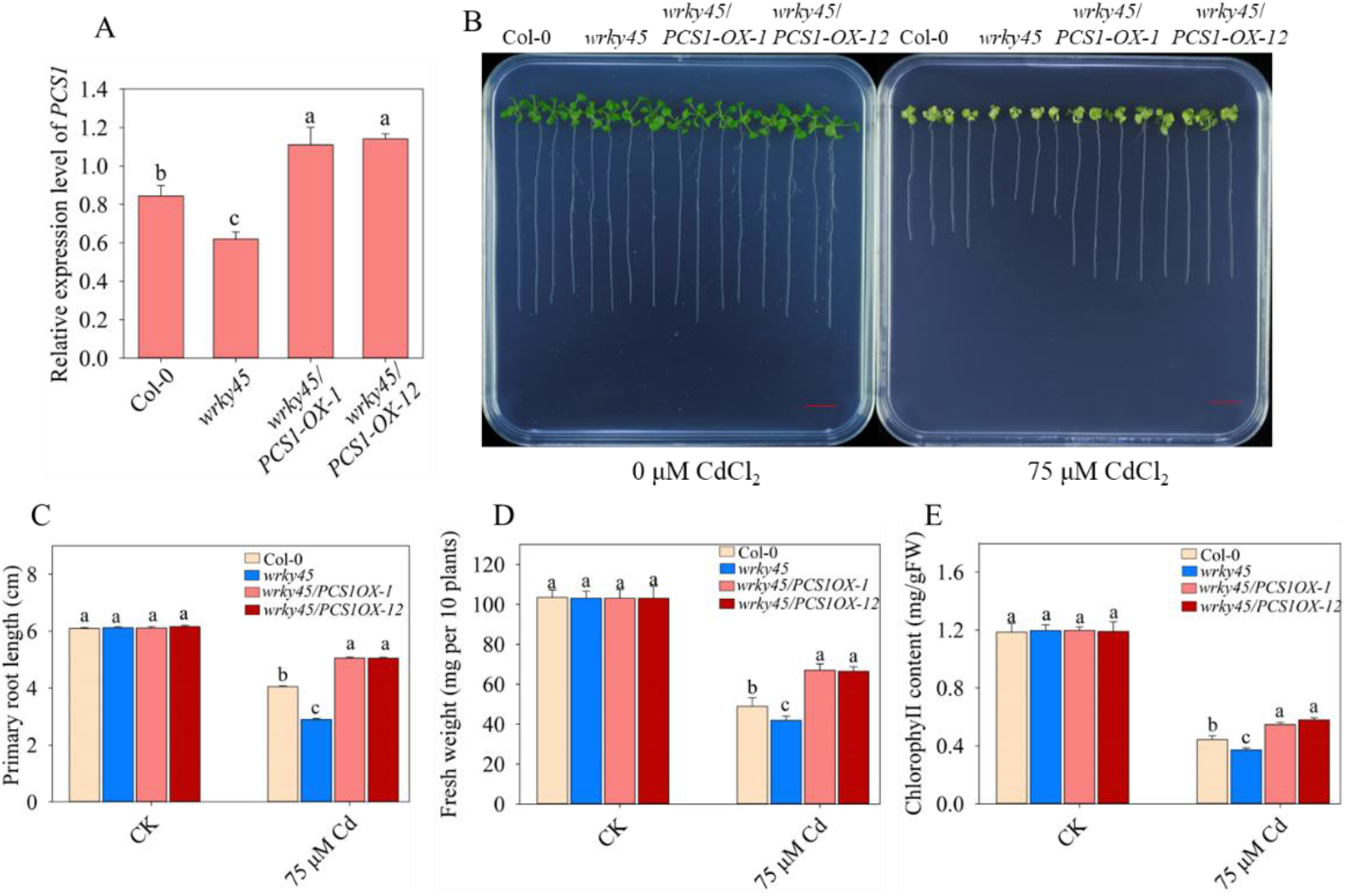
Overexpression of *PCS1* restores the Cd-sensitive phenotype in the *wrky45* mutant. (A) Expression of *PCS1* in *wrky45*, Col-0, and *wrky45/PCS1-OX-1*, *and wrky45/PCS1-OX-1*2 lines. qRT-PCR primers are listed in Supplementary table S1. (B) Growth of Col-0, *wrky45*, *wrky45/PCS1-OX-1*, and *wrky45/PCS1-OX-12* lines under Cd stress. Two-day-old plants grown on half-strength Murashige and Skoog’s (1/2 MS) medium were transferred to 1/2 MS medium without or with 75 μM CdCl_2_. Photographs were taken 10 days after the transfer. Scale bar, 1 cm. (C-E) Primary root length (C), fresh weight (D) and chlorophyll (E) of plants described in (A). Three independent experiments were done with similar results, each with three biologicals repeats. Four plants per genotype from one plate were measured for each repeat. Data are means ± SE (n = 4). Different letters indicate significant difference at P < 0.05 level (one-way ANOVA with Turkey’s test).

**Fig. S5.**
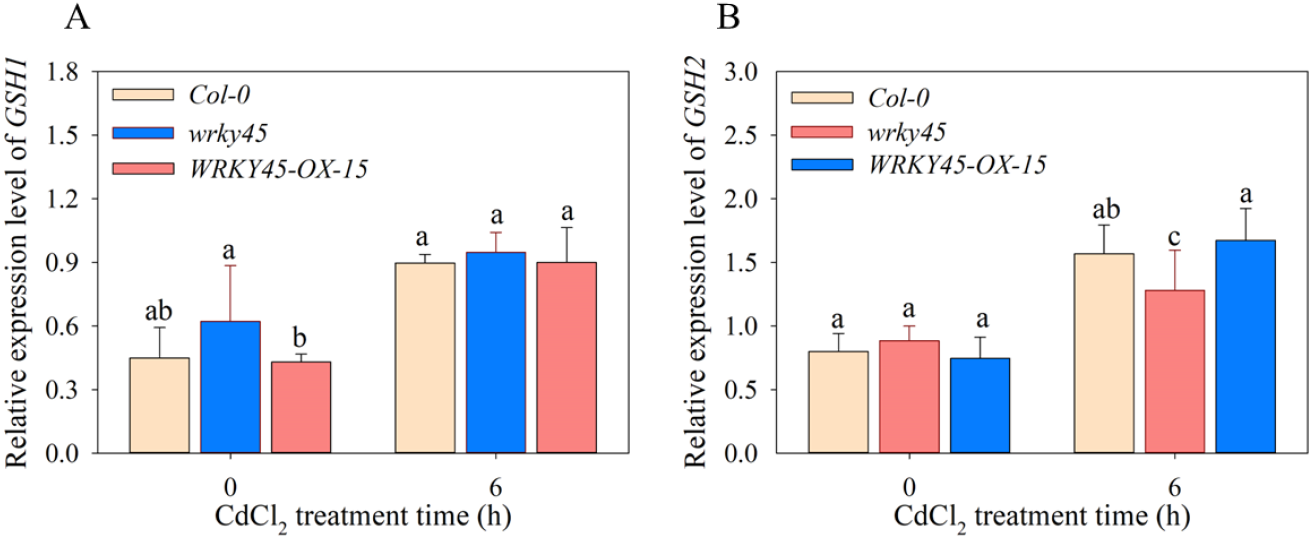
The relative expressions levels of *GSH1*(A), *GSH2* (B) in the wild type (Col-0), *wrky45* mutants, and WRKY45-OX lines revealed by quantitative real time PCR analysis. Three-week-old seedlings were treated with 20 µM CdCl_2_ for 0 h and 6 h. The relative expression was double normalized against *ACTIN2* (*AT3G18780*). Data are means ± SE (n = 4). Different letters indicate significant difference at P < 0.05 level (one-way ANOVA with Turkey’s test).

**Table. S1.**
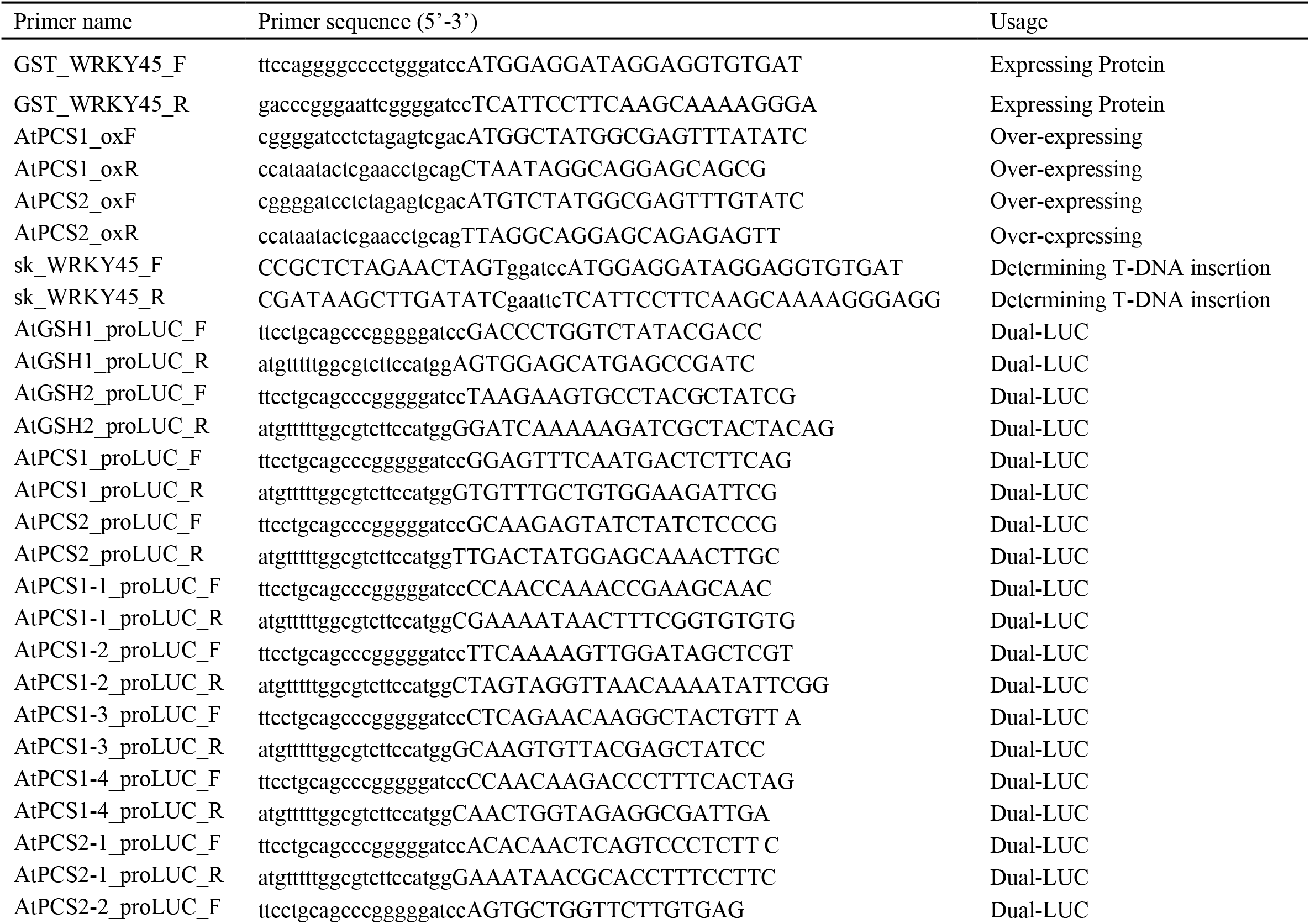
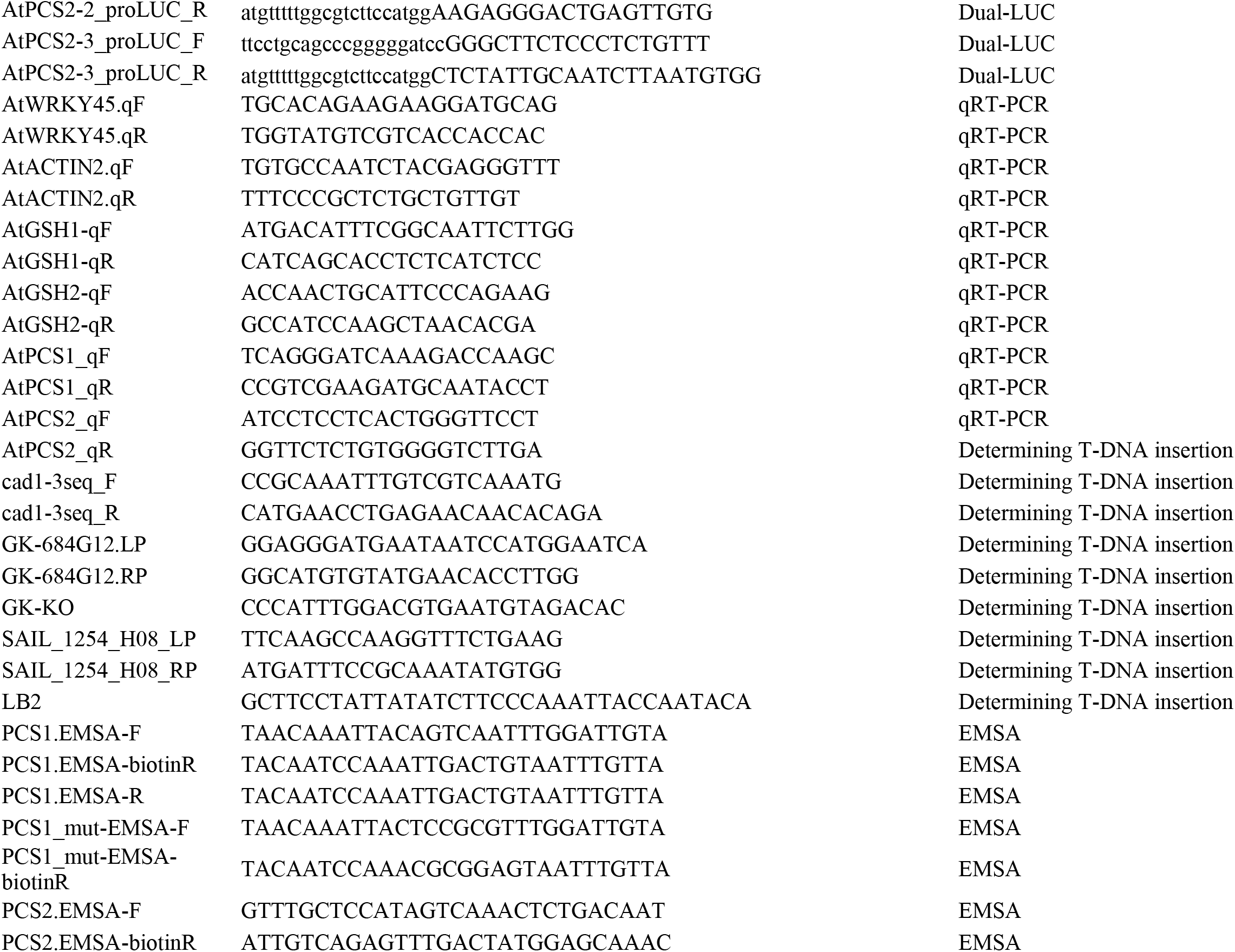
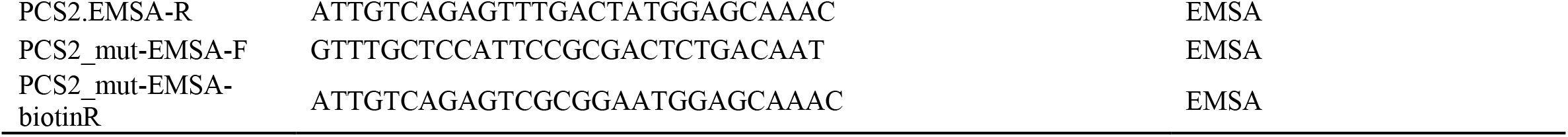
The list of primer pairs used in this study

## References

Agarwal, P., Mitra, M., Banerjee, S., Roy, S., 2020. MYB4 transcription factor, a member of R2R3-subfamily of MYB domain protein, regulates cadmium tolerance via enhanced protection against oxidative damage and increases expression of *PCS1* and *MT1C* in Arabidopsis. Plant Sci, 297,110501.

Andres, S., Andrea, P., 2002. Plant responses to abiotic stresses: heavy metal-induced oxidative stress and protection by mycorrhization. J Exp Bot,53(372),1351–1365.

Barros, J.A.S., Cavalcanti, J.H.F., Pimentel, K.G., Medeiros, D.B., Silva, J.C.F., Condori Apfata, J.A., Lapidot Cohen, T., Brotman, Y., Nunes Nesi, A., Fernie, A.R., Avin Wittenberg, T., Araújo, W.L., 2022. The significance of WRKY45 transcription factor in metabolic adjustments during dark-induced leaf senescence. Plant Cell Environ, 45(9),2682–2695.

Bose, S., Bhattacharyya, A.K., 2008. Heavy metal accumulation in wheat plant grown in soil amended with industrial sludge. Chemosphere,70(7),1264–1272.

Cai Z, Xian P, Wang H, Lin R, Lian T, Cheng Y, Ma Q, Nian H. 2020.Transcription factor GmWRKY142 confers cadmium resistance by up-regulating the Cadmium Tolerance 1-Like genes. Front Plant Sci., 11,724

Chen, J., Nolan, T., Ye, H., Zhang, M., Tong, H., Xin, P., Chu, J., Chu, C., Li, Z., Yin, Y., 2017. Arabidopsis WRKY46, WRKY54 and WRKY70 transcription factors are involved in Brassinosteroid-regulated plant growth and drought response. The Plant Cell, 29(6),1425–1439.

Chen, J., Yang, L., Gu, J., Bai, X., Ren, Y., Fan, T., Han, Y., Jiang, L., Xiao, F., Liu, Y., 2015. MAN3 gene regulates cadmium tolerance through the glutathione-dependent pathway in Arabidopsis thaliana. New Phytol., (2):570–582.

Chen, J., Yang, L., Yan, X., Liu, Y., Wang, R., Fan, T., Ren, Y., Tang, X., Xiao, F., Liu, Y., Cao, S. 2016. Zinc-Finger transcription factor ZAT6 positively regulates cadmium tolerance through the glutathione-dependent pathway in Arabidopsis. Plant Physiol., 171(1),707–719

Chen, L., Xiang, S., Chen, Y., Li, D., Yu, D., 2017. Arabidopsis WRKY45 Interacts with the DELLA protein RGL1 to positively regulate age-triggered leaf senescence. Mol Plant, 10(9),1174–1189.

Ciolkowski, I., Wanke, D., Birkenbihl, R.P., Somssich, I.E. 2008. Studies on DNA-binding selectivity of WRKY transcription factors lend structural clues into WRKY-domain function. Plant Mol Biol., 68(1-2),81–92.

Clemens, S.2001. Molecular mechanisms of plant metal tolerance and homeostasis. Planta, 212(4),475–486.

Clemens, S. 2006. Toxic metal accumulation, responses to exposure and mechanisms of tolerance in plants. Biochimie, 88(11),1707–1719.

Clemens S. 2019. Safer food through plant science: reducing toxic element accumulation in crops. J Exp Bot. 2019, 70(20),5537–5557.

Clough SJ, Bent AF. 1998. Floral dip: a simplified method for Agrobacterium-mediated transformation of Arabidopsis thaliana. Plant J, 6(6),735–43

Cobbett C, Goldsbrough P. 2002.Phytochelatins and metallothioneins: roles in heavy metal detoxification and homeostasis. Annu Rev Plant Biol., 53, 159–182.

Cobbett CS, May MJ, Howden R, Rolls B.1998. The glutathione-deficient, cadmium-sensitive mutant, cad2-1, of Arabidopsis thaliana is deficient in gamma-glutamylcysteine synthetase. Plant J,16(1),73–78.

Cohen CK, Fox TC, Garvin DF, Kochian LV. 1998.The role of iron-deficiency stress responses in stimulating heavy-metal transport in plants. Plant Physiol., 116(3),1063–1072.

DalCorso G, Farinati S, Maistri S, Furini A. 2008. How plants cope with cadmium: staking all on metabolism and gene expression. J Integr Plant Biol.,50(10),1268–1280.

Dang F, Li Y, Wang Y, Lin J, Du S, Liao X. 2022. ZAT10 plays dual roles in cadmium uptake and detoxification in Arabidopsis. Front Plant Sci., 13,994100.

Devaiah BN, Karthikeyan AS, Raghothama KG. 2007. WRKY75 transcription factor is a modulator of phosphate acquisition and root development in Arabidopsis. Plant Physiol., 43(4),1789–801.

Dong J, Chen C, Chen Z. 2003.Expression profiles of the Arabidopsis WRKY gene superfamily during plant defense response. Plant Mol Biol., 51(1),21–37.

Eulgem T, Rushton PJ, Robatzek S, Somssich IE. 2000.The WRKY superfamily of plant transcription factors. Trends Plant Sci., 5(5),199–206.

Fischer S, Kühnlenz T, Thieme M, Schmidt H, Clemens S. 2014. Analysis of plant Pb tolerance at realistic submicromolar concentrations demonstrates the role of phytochelatin synthesis for Pb detoxification. Environ Sci Technol., 48(13),7552–7559.

Fu Y, Li F, Guo S, Zhao M.2021. Cadmium concentration and its typical input and output fluxes in agricultural soil downstream of a heavy metal sewage irrigation area. J Hazard Mater.,412,125203.

Goto S, Sasakura-Shimoda F, Suetsugu M, Selvaraj MG, Hayashi N, Yamazaki M, Ishitani M, Shimono M, Sugano S, Matsushita A, Tanabata T, Takatsuji H. 2015.Development of disease-resistant rice by optimized expression of WRKY45. Plant Biotechnol J., 13(6),753–765.

Grant CA, Clarke JM, Duguid S, Chaney RL. 2008.Selection and breeding of plant cultivars to minimize cadmium accumulation. Sci Total Environ., 390(2-3),301–310.

Han Y, Fan T, Zhu X, Wu X, Ouyang J, Jiang L, Cao S. 2019. WRKY12 represses GSH1 expression to negatively regulate cadmium tolerance in Arabidopsis. Plant Mol Biol., 99(1-2),149–159.

Hong C, Cheng D, Zhang G, Zhu D, Chen Y, Tan M. 2017.The role of ZmWRKY4 in regulating maize antioxidant defense under cadmium stress. Biochem Biophys Res Commun.,482(4),1504–1510.

Howden R, Goldsbrough PB, Andersen CR, Cobbett CS. 1995.Cadmium-sensitive, cad1 mutants of Arabidopsis thaliana are phytochelatin deficient. Plant Physiol.,107(4),1059–1066.

Hu Y, Dong Q, Yu D. 2012.Arabidopsis WRKY46 coordinates with WRKY70 and WRKY53 in basal resistance against pathogen *Pseudomonas syringae*. Plant Sci.,185-186, 288–297.

Huang F, Hu J, Chen L, Wang Z, Sun S, Zhang W, Jiang H, Luo Y, Wang L, Zeng Y, Fang L. 2023. Microplastics may increase the environmental risks of Cd via promoting Cd uptake by plants: A meta-analysis. J Hazard Mater.,448,130887.

Huang, Y.Y., He, C.T., Shen, C., Guo, J.J., Mubeen, S., Yuan, J.G., Yang, Z.Y., 2017.Toxicity of cadmium and its health risks from leafy vegetable consumption. Food Func,1373–1401.

Ichimaru K, Yamaguchi K, Harada K, Nishio Y, Hori M, Ishikawa K, Inoue H, Shigeta S, Inoue K, Shimada K, Yoshimura S, Takeda T, Yamashita E, Fujiwara T, Nakagawa A, Kojima C, Kawasaki T.,2002. Cooperative regulation of PBI1 and MAPKs controls WRKY45 transcription factor in rice immunity. Nat. Commun., 13(1), 2397.

Kahle, H., 1993. Response of roots of trees to heavy metals. Environmental Experimental Botany, 33,99–119.

Khan, I., Asaf, S., Jan, R., Bilal, S., Lubna, Khan, A.L., Kim, K.M., Al-Harrasi, A., 2023. Genome-wide annotation and expression analysis of WRKY and bHLH transcriptional factor families reveal their involvement under cadmium stress in tomato (*Solanum lycopersicum* L.). Front. Plant. Sci.,.14,1100895.

Kim, D., Bovet, L., Maeshima, M., Martinoia, E., Lee, Y., 2007. The ABC transporter AtPDR8 is a cadmium extrusion pump conferring heavy metal resistance. The Plant Journal, 50, 207–218.

Kim, D.Y., Bovet, L., Noh, E.W., Martinoia, E., Lee, Y., 2006. AtATM3 is involved in heavy metal resistance in Arabidopsis. Plant. Physiol., 140(3), 922–932.

Kong, X., Li, C., Zhang, F., Yu, Q., Gao, S., Zhang, M., Tian, H., Zhang, J., Yuan, X., Ding, Z., 2018. Ethylene promotes cadmium-induced root growth inhibition through EIN3 controlled *XTH33* and *LSU1* expression in Arabidopsis. Plant, Cell Environment, 41, 2449–2462.

Korshunova, Y.O., Eide, D., Clark, W.G., Guerinot, M.L., Pakrasi, H.B., 1999. The IRT1 protein from Arabidopsis thaliana is a metal transporter with a broad substrate range. Plant. Mol. Biol.,40,37–44.

Kühnlenz, T., Schmidt, H., Uraguchi, S., Clemens, S., 2014. Arabidopsis thaliana phytochelatin synthase 2 is constitutively active in vivo and can rescue the growth defect of the PCS1-deficient *cad1-3* mutant on Cd-contaminated soil. J. Exp. Bot, 4241-4253.

Lee S, Kim YY, Lee Y, An G.,2007. Rice P1B-type heavy-metal ATPase, OsHMA9, is a metal efflux protein. Plant.Physiol.,145(3):831–842.

Li GZ, Zheng YX, Liu HT, Liu J, Kang GZ.,2022.WRKY74 regulates cadmium tolerance through glutathione-dependent pathway in wheat. Environ. Sci. Pollut. Res. Int.,29(45):68191–68201.

Li, J., Besseau, S., Toronen, P., Sipari, N., Kollist, H., Holm, L., Palva, E.T., 2013. Defense-related transcription factors WRKY70 and WRKY54 modulate osmotic stress tolerance by regulating stomatal aperture in Arabidopsis. New. Phytol., 200, 457–472.

Li, S., Fu, Q., Chen, L., Huang, W., Yu, D., 2011. Arabidopsis thaliana WRKY25, WRKY26, and WRKY33 coordinate induction of plant thermotolerance. Planta, 233(6),1237–1252.

Li, W., Wang, H., Yu, D., 2016. Arabidopsis WRKY transcription factors WRKY12 and WRKY13 oppositely regulate flowering under short-day conditions. Mol. Plant., 9(11):1492–1503.

Liu, H., Yu, X., Li, K., Klejnot, J., Yang, H., Lisiero, D., Lin, C., 2008. Photoexcited CRY2 Interacts with CIB1 to Regulate Transcription and Floral Initiation in Arabidopsis. Science, 322,1535–1539.

Liu Y, He G, He Y, Tang Y, Zhao F, He T., 2023. Discovery of cadmium-tolerant biomacromolecule (StCAX1/4 transportproteins) in potato and its potential regulatory relationship with WRKY transcription factors. Int. J. Biol. Macromol., 228:385–399.

López-Millán, A., Sagardoy, R., Solanas, M., Abadía, A., Abadía, J., 2009. Cadmium toxicity in tomato (Lycopersicon esculentum) plants grown in hydroponics. Environmental. Experimental. Botany., 65,376–385.

Mendoza-Cózatl, D., Loza-Tavera, H., Hernández-Navarro, A., Moreno-Sánchez, R., 2005. Sulfur assimilation and glutathione metabolism under cadmium stress in yeast, protists and plants. FEMS. Microbiol. Rev., 29(4),653–671.

Mendoza-Cozatl, D.G., Jobe, T.O., Hauser, F., Schroeder, J.I., 2011. Long-distance transport, vacuolar sequestration, tolerance, and transcriptional responses induced by cadmium and arsenic. Curr. Opin. Plant. Biol,14,554–562.

Meng, X., Li, W., Shen, R., Lan, P., 2022. Ectopic expression of IMA small peptide genes confers tolerance to cadmium stress in Arabidopsis through activating the iron deficiency response. J. Hazard. Mater, 422,126913.

Nam, H., Shahzad, Z., Dorone, Y., Clowez, S., Zhao, K., Bouain, N., Lay-Pruitt, K.S., Cho, H., Rhee, S.Y., Rouached, H., 2021. Interdependent iron and phosphorus availability controls photosynthesis through retrograde signaling. Nat. Commun., 12(1),7211.

Nazar, R., Iqbal, N., Masood, A., Khan, M.I.R., Syeed, S., Khan, N.A., 2012. Cadmium Toxicity in Plants and Role of Mineral Nutrients in Its Alleviation. American Journal of Plant Sciences, 03,1476–1489.

Ou, S., Xu, Z., Mai, C., Li, B., Wang, J., 2022. Ectopic expression of GmNF-YA8 in Arabidopsis delays flowering via modulating the expression of gibberellic acid biosynthesis-and flowering-related genes and promotes lateral root emergence in low phosphorus conditions. Front. Plant.Sci.,13, 1033938

Peng, J., Wang, Y., Ding, G., Ma, H., Zhang, Y., Gong, J., 2017. A Pivotal Role of Cell Wall in Cadmium Accumulation in the Crassulaceae hyperaccumulator Sedum plumbizincicola. Mol. Plant., 10,771–774.

Reeves PG, Chaney RL., 2008. Bioavailability as an issue in risk assessment and management of food cadmium: a review. Sci. Total. Environ., 398(1-3),13–19.

Riaz M, Kamran M, Rizwan M, Ali S, Parveen A, Malik Z, Wang X., 2021. Cadmium uptake and translocation: selenium and silicon roles in Cd detoxification for the production of low Cd crops: a critical review. Chemosphere., 273,129690.

Rushton, P.J., Somssich, I.E., Ringler, P., Shen, Q.J., 2010. WRKY transcription factors. TRENDS PLANT SCI15,247–258.

Scarpeci, T.E., Zanor, M.I., Mueller-Roeber, B., Valle, E.M., 2013. Overexpression of *AtWRKY30* enhances abiotic stress tolerance during early growth stages in Arabidopsis thaliana. Plant. Mol. Biol., 83,265–277.

Shen, N., Hou, S., Tu, G., Lan, W., Jing, Y., 2021. Transcription Factor WRKY33 mediates the phosphate deficiency-induced remodeling of root architecture by modulating iron homeostasis in Arabidopsis roots. International Journal of Molecular Sciences, 22,9275.

Sheng, Y., Yan, X., Huang, Y., Han, Y., Zhang, C., Ren, Y., Fan, T., Xiao, F., Liu, Y., Cao, S., 2019. The WRKY transcription factor, WRKY13, activates PDR8 expression to positively regulate cadmium tolerance in Arabidopsis. Plant. Cell. Environ.,42,891–903.

Shimono M, Koga H, Akagi A, Hayashi N, Goto S, Sawada M, Kurihara T, Matsushita A, Sugano S, Jiang CJ, Kaku H, Inoue H, Takatsuji H., 2012.Rice WRKY45 plays important roles in fungal and bacterial disease resistance. Mol. Plant. Pathol., 13(1):83–94.

Su, T., Xu, Q., Zhang, F., Chen, Y., Li, L., Wu, W., Chen, Y., 2015. WRKY42 Modulates Phosphate Homeostasis through Regulating Phosphate Translocation and Acquisition in Arabidopsis. Plant Physiol.,167,1579–1591.

Sun, Y., Liu, Z., Guo, J., Zhu, Z., Sun, X., 2020. WRKY33-PIF4 loop is required for the regulation of H2O2 homeostasis. Biochem. Biophys. Res. Commun., 527(4),922–928.

Tang, J., Wu, D., Li, X., Wang, L., Xu, L., Zhang, Y., Xu, F., Liu, H., Xie, Q., Dai, S., Coleman-Derr, D., Zhu, S., Yu, F., 2022. Plant immunity suppression via PHR1-RALF-FERONIA shapes the root microbiome to alleviate phosphate starvation. EMBO J, 41(6): e109102.

Tang, R., Yang, Y., Yan, Y., Mao, D., Yuan, H., Wang, C., Zhao, F., Luan, S., 2022. Two transporters mobilize magnesium from vacuolar stores to enable plant acclimation to magnesium deficiency. Plant Physiol.,190,1307–1320.

Tang, Z., Wang, H.Q., Chen, J., Chang, J.D., Zhao, F.J., 2023. Molecular mechanisms underlying the toxicity and detoxification of trace metals and metalloids in plants. J Integr Plant Biol.,65(2),570–593.

Tong Y, Gao J, Yue T, Zhang X, Liu J, Bai J.,2023, Distribution, chemical fractionation, and potential environmental risks of Hg, Cr, Cd, Pb, and as in wastes from ultra-low emission coal-fired industrial boilers in China. J. Hazard. Mater., 446:130606.

Ulker, B., Somssich, I.E., 2004.WRKY transcription factors: from DNA binding towards biological function. Curr. Opin.Plant. Biol., 7, 491–498.

Wang, H., Xu, Q., Kong, Y., Chen, Y., Duan, J., Wu, W., Chen, Y., 2014. Arabidopsis WRKY45 Transcription factor activates PHOSPHATE TRANSPORTER1;1 expression in response to phosphate starvation. Plant Physiology,164,2020–2029.

Wang, P., Chen, H., Kopittke, P.M., Zhao, F., 2019. Cadmium contamination in agricultural soils of China and the impact on food safety. Environmental Pollu.,249,1038–1048.

Wang, Y., Schuck, S., Wu, J., Yang, P., Döring, A., Zeier, J., Tsuda, K., 2018. A MPK3/6-WRKY33-ALD1-Pipecolic Acid Regulatory Loop Contributes to Systemic Acquired Resistance. The Plant Cell, 30, 2480–2494.

Wei, W., Zhang, Y., Han, L., Guan, Z., Chai, T., 2008. A novel WRKY transcriptional factor from Thlaspi caerulescens negatively regulates the osmotic stress tolerance of transgenic tobacco. Plant Cell Reports, 27,795–803.

Wu, H., Chen, C., Du, J., Liu, H., Cui, Y., 2012. Co-overexpression FIT with AtbHLH38 or AtbHLH39 in Arabidopsis-enhanced cadmium tolerance via increased cadmium sequestration in roots and improved iron homeostasis of shoots. Plant Phy., 158,790–800.

Wu, S., Yang, Y., Qin, Y., Deng, X., Zhang, Q., Zou, D., Zeng, Q., 2023. Cichorium intybus L. is a potential Cd-accumulator for phytoremediation of agricultural soil with strong tolerance and detoxification to Cd. J. Hazard. Mater.,451,131182.

Wu, X., Chen, L., Lin, X., Chen, X., Han, C., Tian, F., Wan, X., Liu, Q., He, F., Chen, L., Zhong, Y., Yang, H., Zhang, F., 2023. Integrating physiological and transcriptome analyses clarified the molecular regulation mechanism of PyWRKY48 in poplar under cadmium stress. Int. J. Biol. Macromol., 238,124072.

Wu, X., Chen, Q., Chen, L., Tian, F., Chen, X., Han, C., Mi, J., Lin, X., Wan, X., Jiang, B., Liu, Q., He, F., Chen, L., Zhang, F., 2022. A WRKY transcription factor, PyWRKY75, enhanced cadmium accumulation and tolerance in poplar. Ecotox. Environ. Safe,239,113630.

Xian, P., Yang, Y., Xiong, C., Guo, Z., Alam, I., He, Z., Zhang, Y., Cai, Z., Nian, H., 2023. Overexpression of GmWRKY172 enhances cadmium tolerance in plants and reduces cadmium accumulation in soybean seeds. Front. Plant Sci.,14, 1133892.

Yang, G., Wang, C., Wang, Y., Guo, Y., Zhao, Y., Yang, C., Gao, C., 2016. Overexpression of ThVHAc1 and its potential upstream regulator, ThWRKY7, improved plant tolerance of Cadmium stress. Sci. Repor.,6, 18752.

Zhang, C., Tong, C., Cao, L., Zheng, P., Tang, X., Wang, L., Miao, M., Liu, Y., Cao, S., 2023. Regulatory module WRKY33-ATL31-IRT1 mediates cadmium tolerance in Arabidopsis. Plant Cell Environment, 46,1653–1670.

Zhang, H., Zhang, L., Ji, Y., Jing, Y., Li, L., Chen, Y., Wang, R., Zhang, H., Yu, D., Chen, L., 2022. Arabidopsis SIGMA FACTOR BINDING PROTEIN1 (SIB1) and SIB2 inhibit WRKY75 function in abscisic acid-mediated leaf senescence and seed germination. J. Exp. Bot.,73,182–196.

Zhang, L., Chen, L., Yu, D., 2018. Transcription Factor WRKY75 Interacts with DELLA Proteins to Affect Flowering. Plant Physiology,176,790–803.

Zhang, Q., Cai, W., Ji, T., Ye, L., Lu, Y., Yuan, T., 2020. WRKY13 enhances cadmium tolerance by promoting D-CYSTEINE DESULFHYDRASE and hydrogen sulfide production. Plant Physiology,183,345–357.

Zhao, F.J., Tang, Z., Song, J.J., Huang, X.Y., Wang, P., 2022. Toxic metals and metalloids: uptake, transport, detoxification, phytoremediation, and crop improvement for safer food. Mol. Plant.,15,27–44.

Zhao, M., Wang, H., Sun, J., Tang, R., Cai, B., Song, X., Huang, X., Huang, J., Fan, Z., 2023. Spatio-temporal characteristics of soil Cd pollution and its influencing factors: A Geographically and temporally weighted regression (GTWR) method. J. Hazard. Mater,446,130613.

Zheng, Z., Qamar, S.A., Chen, Z., Mengiste, T., 2006. Arabidopsis WRKY33 transcription factor is required for resistance to necrotrophic fungal pathogens. The Plant Journal,48,592–605.

Zhu, S., Chen, M., Liang, C., Xue, Y., Lin, S., Tian, J., 2020. Characterization of purple acid phosphatase family and functional analysis of GmPAP7a/7b involved in extracellular ATP utilization in soybean. Front. Plant. Sci.,11,661.

Zhu, Y.X., Du, W.X., Fang, X.Z., Zhang, L.L., Jin, C.W., 2020. Knockdown of BTS may provide a new strategy to improve cadmium-phytoremediation efficiency by improving iron status in plants. J. Hazard. Mater,84,121473.

